# Realistic alpha oscillations and transient responses in a cortical microcircuit model

**DOI:** 10.1101/2021.11.10.468067

**Authors:** Andres A. Kiani, Theoden I. Netoff, Geoffrey M. Ghose

## Abstract

Neural-mass modeling of neural population data (EEG, ECoG, or LFPs) has shown promise both in elucidating the neural processes underlying cortical rhythms and changes in brain state, as well as offering a framework for testing the interplay between these rhythms and information processing. Models of cortical alpha rhythms (8 - 12 Hz) and their impact in visual sensory processing have been at the forefront of this effort, with the Jansen-Rit being one of the more popular models in this domain. The Jansen-Rit model, however, fails in reproducing key physiological observations including the level of inputs that cortical neurons receive and their responses to visual transients. To address these issues we generated a neural mass model that complies better with synaptic mediated dynamics, intrinsic alpha behavior, and produces realistic responses. The model is robust to many changes in parameter values but critically depends on the ratio of excitation to inhibition, producing response transients whose features are dependent on this ratio and alpha phase and power. The model is sufficiently flexible so as to be able to easily replicate the range of low frequency oscillations observed in different studies. Consistent with experimental observations, we find phase-dependent response dynamics to both visual and electrical stimulation using this model. The model suggests that stimulation facilitates alpha at particular phases and suppresses it in others due to a phase dependent lag in inhibitory responses. Hence, the model generates insight into the physiological parameters responsible for intrinsic oscillations and testable hypotheses regarding the interactions between visual and electrical stimulation on those oscillations.

**Author summary:** While there is increasing evidence of the fundamental role brain states play in shaping information processing in the cerebral cortex, a mechanistic understanding of how those brain states are manifested and alter the signals underlying sensory processing and decision making has proved challenging. To address this issue we have modeled a well established signature of inattention in visual cortex: synchronized low frequency (8 - 12 Hz) oscillations. To allow for inferences regarding the local generation of these rhythms within a cortical microcircuit we used a neural mass model approach that incorporates physiologically realistic interactions between 3 populations of neurons. Our model is able to explain a variety of experimental observations that previous neural mass models have not, including spontaneous rhythms in the absence of input, the faithful transmission of strong input transients, a range of oscillation frequencies, and phase dependent visual responses. The model is robust to a range of parameters, but critically depends on the balance between local excitation and inhibition.

## Introduction

Many neural recording techniques, such as electroencephalograms (EEG), electrocorticogram (ECoG), and local field potentials (LFPs), record population-level activity that directly reflects information processing and behavioral state. These signals can exhibit complex spatiotemporal patterns in response to stimuli, cognitive state, and behavioral changes. [67] An understanding of the underlying dynamics producing these complicated signals is further complicated by the complex nature of the biological neural networks implicated in their production, with diverse classes of neurons whose interactions are mediated by the sophisticated interplay between channel, membrane, and synaptic dynamics. [32] The difficulty of the problem is moreover amplified by the interaction of these measured signals with non-neuronal tissues. [13]. Consequently, uncovering the underlying processes associated with these signals has been an immense challenge.

In view of these difficulties, researchers have turned toward the development of computational models that are able to capture the overlying dynamics including various phenomena such as the alpha rhythms of the visual cortex, whilst being capable of making inferences regarding underlying processes like excitation-inhibition balance. [23] Various modeling approaches have been endeavored in this domain, out of which the Neural-mass modeling approach has become widely popularized in the literature. [66] “Neural-mass” or “Mean-field” computational models, first introduced by Wilson and Cowan, adopt a reductionist approach by modeling the activity of interconnected populations of inhibitory and excitatory cells through coupled non-linear differential equations. [14] Such simplified models, when they include non-linear and delayed interactions between idealized sub-populations of neurons, can successfully model the dynamics of local field potentials. [61]

For example, Lopes da Silva et al., proposed one of the earliest models using coupled populations of excitatory and inhibitory neurons to simulate the generation of alpha rhythms (8 - 12 Hz), long associated with inattention in EEG studies. [59, 53] Jansen and Rit extended this model by adding a pyramidal population to explain how such rhythms within a local cortical population of the visual cortex are generated and how the circuit is affected by incoming signals. [29, 31]

Although the Jansen-Rit (JR) model successfully models how alpha rhythms are modulated in a phase dependent manner by stimulation, it does not accurately simulate several aspects of cortical responses.[41] First, alpha rhythms are largely associated with relative quiescence [1, 15, 36]: the first observations of cortical alpha were obtained from subjects with their eyes closed in the absence of any visual stimulation.[6] However, stable oscillations in the absence of parameter manipulations emerges in the JR system when driven by drive abpve 155 Hz. [23, 30] Second, the stable oscillations come at the cost of realistic response dynamics to input transients, which are largely lost, or substantially altered, in the output of these models [41, 17]. For example, stable oscillations in the JR system are seen at input rates as high as 315 Hz, precluding the faithful transmission of high input transients.

A related issue is the mismatch between received input and output rates, which is a necessary self-consistency constraint given the interconnectedness of cortical columns both within and between areas. Cortical regions have roughly the same structure, consist of similar neuronal species, and exhibit similar dynamics at a local level. [21] While primary visual cortex (V1) receives primary input from thalamic neurons, all other visual areas, including V2, V4, MT, and IT, receive their input from other cortical areas [27, 47]. Therefore, input and output rates should be comparable, if not the same, in an adequate cortical model. As it stands, the upper bound imposed on a population output in the Jansen-Rit is 5 Hz. The average maximum single unit firing rate in both afferent V1 and efferent V4 neurons is around or below 50 Hz to a preferred stimuli [45, 65]. Therefore, it stands to reason that the average maximum output rate across a cortical population (representing thousands or millions of neurons) during a visually evoked response should be some factor lower than that of the average single unit due to the vast diversity of feature and spatial preferences represented across the ensemble.

Another critical flaw in the Jansen-Rit model is the linear mapping between input rates and synaptic drives, a clear contradiction from synaptic mediated dynamics and inconsistent with non-linearities observed in visual receptive fields [57]. The linear relationship between input rate and synaptic drive in potential results in linear response dynamics that do not include thresholds, saturations, and attenuations, all of which are dynamical features observed in biological neural networks. The nonlinear nature of membranes and synapses is responsible for the emergence of multiple stable states and the ability to transition between distinct modes of response, all of which are hindered in this linear regime.

A final concern is related to the Jansen-Rit’s inability to generate the different frequencies observed in neural recording data, even though the population time constants of the model were intended to serve this purpose. Previous studies have either introduced additional populations, or additional connections with associated connectivity factors, as an attempt to resolve this problem.

In this manuscript we show that simple physiologically informed modifications of the classic three population model can address all of these limitations. Like the Jansen-Rit model, this new model exhibits alpha rhythm activity that can be disrupted by input; however, in this model the rhythms are only prevalent when inputs are relatively inactive and are notably absent in the regime of strong input, allowing for a faithful transmission of input transients. The model is self consistent, in that input and output rates are comparable, and incorporates the fact that the effects of external drive on membrane potentials are synaptically mediated. Additionally, we formulate a relationship between the model’s time constants and synaptic capacitances that effectively resolves the frequency problem, allowing the three population model to exhibit the myriad of physiologically relevant frequencies. Finally, because oscillations are only observed in particular coupling regimes, the model suggests that cortical function is strongly dependent on the maintenance of excitatory-inhibitory coupling ratios over a limited range.

## Results

In this paper we introduce the self-consistent intrinsically oscillating microcircuit (SCIOM) model. The model is an extension of the Janset-Rit three population model of cortical microcircuitry (Fig. 1) with a combination of structural and parameter changes to create a more physiologically realistic model that is able to explain critical experimental observations. Unlike the classic Jansen-Rit model, it produces intrinsic alpha rhythms that are suppressed with increased external inputs. Unlike the Jansen-Rit, there are both physiological bounds on inputs and consistency between input and output rates. And unlike the Jansen-Rit, responses to inputs are mediated by nonlinearities akin to the synaptic nature of biological neural networks.

**Fig 1.**
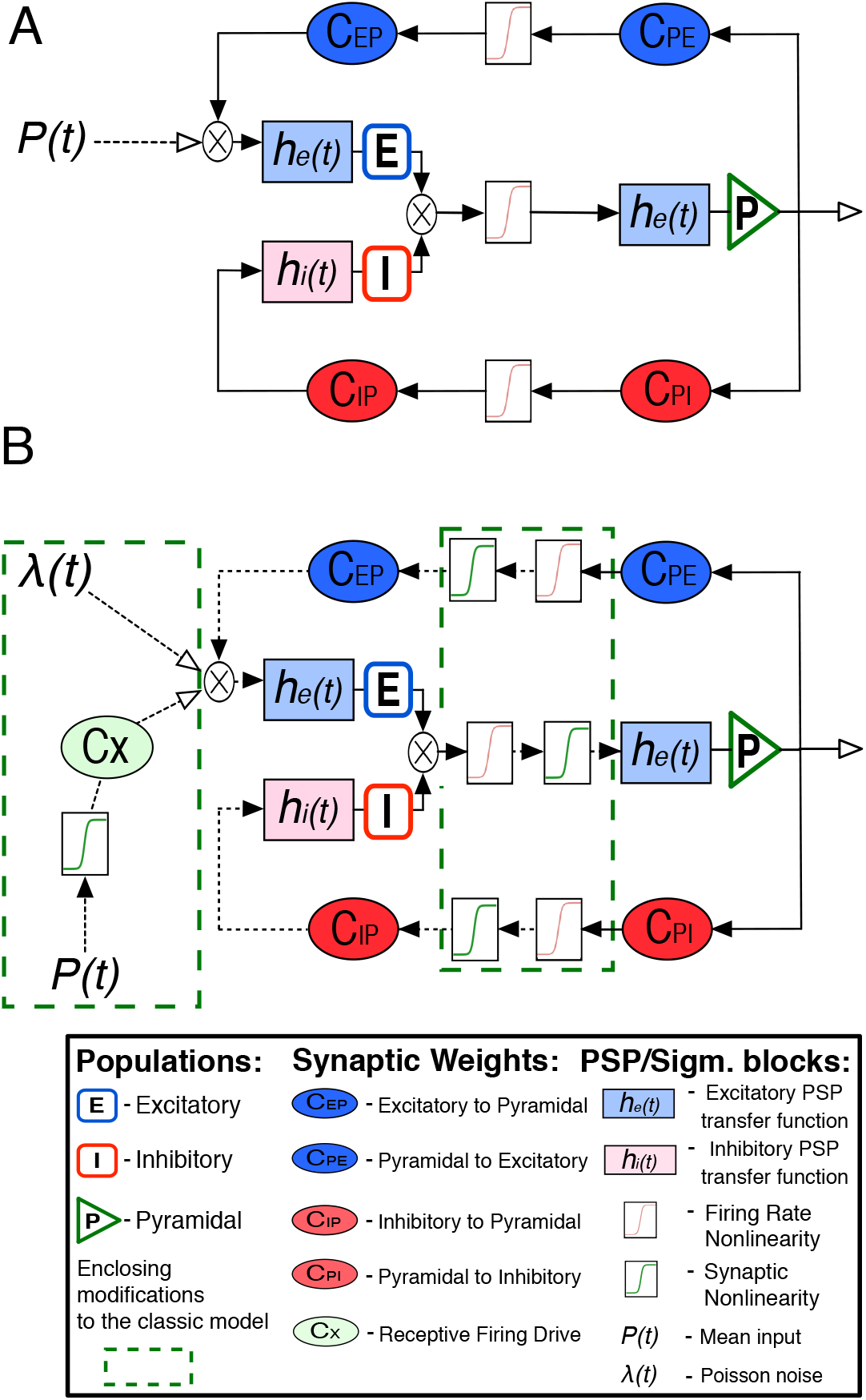
Block diagram schematic of the Jansen-Rit model (A) and the SCIOM model (B). New components added to the Jansen-Rit model to create the SCIOM model are shown in the green box in panel B. The connection between populations is shown as a sequence of arrows. These arrows depict the drive from the presynaptic population to the postsynaptic population. Dashed arrows indicate drive in terms of rate, while solid arrows indicate drive in terms of electric potential. Blocks indicate transformation for synaptic and sigmoidal functions. Ellipses indicate coupling coefficients that linearly scale drive. Stadiums (interneurons) and triangles (pyramidal) symbolize the populations in terms of their second order membrane potential function. The operator symbol indicates the superposition of drives from afferent sources.

First, we will examine several representative equilibrium curves of the Jansen-Rit and the SCIOM systems that illuminate important features and highlight key differences and similarities between the two models, with particular focus on the presence of spontaneous oscillations in the absence of external drive and their suppression with increased drive. We will then adopt an energy approach to gain insight into the parameters and features of the SCIOM model responsible for the emergence of intrinsic oscillations and the interactions of these oscillations with visually evoked transients. We use this energy approach to map how the power and frequency of those oscillations depends on particular parameters.

### Equilibria and State Transitions

The first experimental observation we focus on is the presence of intrinsic oscillations, which occur in the absence of significant input and whose magnitude decreases with strong inputs so as to allow such inputs to propagate. Both of these observations are consistent with experimental observations in visual cortex. We therefore derive equilibrium curves of the local field potential, *Y*, as a function of external input levels, *P* (Eq. 25). Consistent with previous work, and experimental data demonstrating the dominance of synaptic potentials in local field potentials, we define this *Y* as the difference between the excitatory and inhibitory membrane potentials. Doing so allows one to partition equilibrium curves into four segments: the high and low fixed point segments (**Fix_L_** and **Fix_H_**), a separatrix (**SP**), and the stable limit-cycle segment (**LC**).

In Fig. 2A we see that the standard parameter Janset-Rit equilibrium curve’s **LC** domain does not include *P* = 0 (i.e., it does not intrinsically oscillate), but rather stable oscillations are only observed with high levels of input drive. Furthermore, within the first half-domain of **LC**, increasing *P* results in the strengthening of the system’s oscillatory power (*∂*|Ψ| > 0), indicated by the green shading. The waxing and waning of alpha power with increasing *P* across the two aforementioned subsegments of **LC** is observed in simulations of the standard system, shown in the inset. Moreover, the end of **LC** occurs at *Y*(*P* = 320 Hz) = 8.00 mV, which means that any external input function *P*(*t*) ≤ 320 Hz will be effectively attenuated by the system.

**Fig 2.**
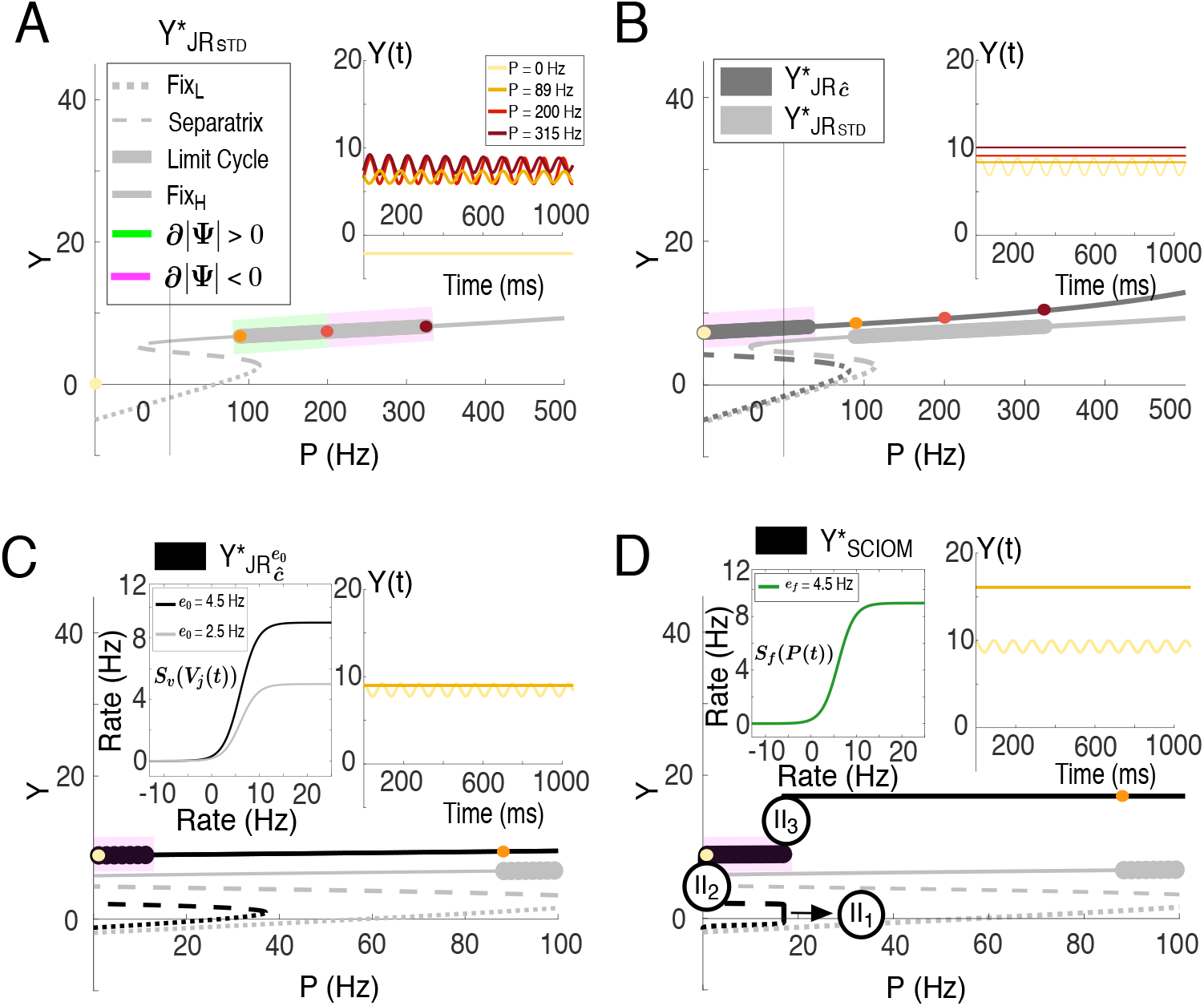
Equilibrium curve transition from the standard Jansen-Rit model to the SCIOM model. The standard Jansen-Rit’s equilibrium curve, 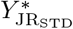, as a function of external input, *P*. The curve is divided into the four segments: low amplitude fixed point, **Fix_L_**, the bistable region **SP**, the stable limit-cycle **LC**, and high amplitude fixed point **Fix_H_**. The curve exhibits an initial Hopf-Bifurcation *Y*(*P* = 89) = 6.74 and then a subsequent Hopf-Bifurcation at *Y*(*P* = 320) = 8.10, defining the boundary of the limit-cycle. The **LC** is further divided into two segments according to the relationship between oscillatory power (|Ψ|) and external drive, *P*. The green shading indicates a positive relationship between |Ψ| and *P* such that the oscillation grows with increasing *P*, while the pink segment indicates a negative one, where the oscillation decreases with increasing *P*. The inset shows time-series corresponding to points on the the equilibrium curve with constant input-rates of *P* ≡ 0, 89, 200, and 320 Hz. (B) The parameter adapted Jansen-Rit model, 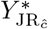 exhibits an initial Hopf-Bifurcation *Y*(*P* =−34) = 6.56 and a subsequent Hopf-Bifurcation *Y*(*P* = 45) = 7.20. For reference, 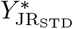 is plotted in gray. (C) The parameter adapted Jansen-Rit model, 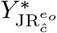, with an adapted *S_F_* nonlinearity in black, plotted over the standard *S*_F_ nonlinearity in grey. Note that the input drive range in this panel has been narrowed to the range of physiological drive. (D) Presents a SCIOM equilibrium curve, 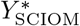. Note the stepwise structure between the partitions: **Fix_L_** and **LC**, divided by steps: **II**_1_, **II**_2_, and **II**_3_ where the system evolves independently of *P*.

These failures to reproduce appropriate oscillatory or limit cycle behavior led us to alter the coupling parameters of the original Jansen-Rit model to investigate how intrinsic oscillations that decay with strong input might be created.

With respect to inhibition, we found that increases in either of the inhibitory coupling weights (the inhibitory to pyramidal connection and the pyramidal to inhibitory connections: *C*_IP_ and *C*_PI_) produce a positive translation in the **LC** segment about the *P* coordinate of the *Y* vs. *P* phase-space portrait, implying that greater external drive, *P*, is required for producing stable oscillations. Thus there is an upper bound, 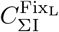, for net inhibition beyond which intrinsic oscillations (*P* = 0) do not occur and the system inevitably converges to **Fix_L_**. Increased inhibition also increases the range over which **LC** is observed and there is therefore a lower bound, 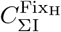, for net inhibition, below which **LC** is not observed and the system inevitably converges to **Fix_H_**.

As would be expected given the definition of LFP signals as the difference between excitatory and inhibitory potentials, changes in excitation have largely opposite effects than to inhibition. Increases in either of the excitatory coupling weights (the excitatory to pyramidal connection and the pyramidal to excitation connection: *C*_EP_ and *C*_PE_), produce a negative translation in the **LC** segment about the *P* coordinate of the *Y* vs. *P* phase-space portrait, suggesting that less external drive, *P*, is required for producing stable oscillations. Thus there is a lower bound, 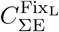, for net excitation beyond which intrinsic oscillations (*P* = 0) do not occur and the system converges to **Fix_L_**. Decreased excitation also increases the range over which **LC** is observed and there is therefore an upper bound, 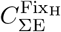, for net excitation, above which **LC** is not observed and the system converges to **Fix_H_**.

Irrespective of class (excitatory or inhibitory), there is a slight asymmetry in the impact that pyramidal to interneuronal couplings have on **LC** because the firing-rate nonlinearity, *S*_F_, operates on *C*_PE_ and *C*_PI_, but not on *C*_EP_ and *C*_IP_ in the Jansen-Rit formulation. Specifically, because |*∂S_F_*| > 1 for most of the responsive domain of *S*_F_, *C*_PE_ and *C*_PI_ are more sensitive than *C*_EP_ and *C*_IP_. Note that this is not the case in standard parameter Jansen-Rit.

These observations characterize the dependency of the system’s oscillatory behavior on the coupling between the three populations. They were utilized to derive coupling combinations that result in intrinsic oscillations that decay in power with the introduction and intensification of input and that disappear within *P* ≤ 2*e_o_*, where 2*e_o_* is the upper bound of, *S_F_*.

The equilibrium curve in Fig. 2B illustrates the common dynamical features possessed by Jansen-Rit coupling configurations that support: intrinsic oscillations, as indicated by the intersection of **LC** and the vertical axis at *P* = 0); their decay (*∂*|Ψ| < 0) with increased input, indicated by the pink shading across the positive *P* domain of LC; and their dissolution within the bounds of the firing-rate nonlinearity, *S_F_*, illustrated by the termination of the **LC** segment within *P* ≤ 2*e_o_*.

Although there are many integer-valued coupling configurations with identical net excitation and inhibition coupling ratios (perhaps infinitely many) that satisfy these three conditions on **LC**, there are very few coupling configurations with different integer-valued coupling ratios that do. Choosing to raise *S*_F_’s upper bound from the 2*e_o_* = 5 Hz to 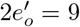 Hz so that outputs better reflect the magnitude and dynamics of inputs, also increased the range of excitatory-inhibitory coupling ratios that result in intrinsic oscillatory behavior. The left inset in Fig. 2C compares the graph of *S*_F_ with an upper bound of 5 Hz to that with an upper bound of 10 Hz. And in Fig. 2C we produce an example equilibrium curve of parameter modified Jansen-Rit systems with this heightened *S*_F_ that satisfy the conditions we have set for intrinsic oscillatory behavior.

Even though a number of conditions are satisfied with these parameter modifications, as we will discuss later, they are not sufficient to replicate the dynamics of realistic response transients. In particular, such systems are not capable of producing nonlinear transformations of input that are physiologically appropriate.

Therefore, we derive equilibrium curves of the local field potential, *Y*, as a function of external input levels, *P*, for the SCIOM model (Eq. 36). Partitioning SCIOM equilibrium curves in terms of the four equivalence relations: **Fix_L_**, **SP**, **LC**, **Fix_H_**; highlights several distinctive features relevant to the SCIOM family of equilibrium curves not seen in Jansen-Rit, as is shown in Fig. 2D. Particularly apparent are the step-wise divisions between the input-dependent segments of *Y*_SCIOM_. We mark these input-independent segments as: **II**_1_, **II**_2_, and **II**_3_. Such input-independence is not observed in the Jansen-Rit model, which is influenced by external inputs across the entire domain of *P*, even when the system is non-responsive to network interactions (see Appendix D for more details). This behavior is responsible for the Jansen-Rit model’s unbounded replication of inputs, a point that we will investigate further in later result sections.

By contrast in SCIOM, segments **II**_1_ and **II**_3_ are associated with the saturation of the system’s response to extra-network drive. Given the parameter values in Table 2, this saturation point occurs at *P* ≤17 Hz, while the segment **II**_2_ is associated with the attenuation of extra-network drive, for *P* ≤2 Hz. See Appendix D for a more rigorous analysis of these features and saturation points.

In the SCIOM paradigm, the transition between **Fix_L_** and **SP** happens at inputs lower than or equal to the maximal firing-rate upperbound 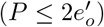, as *P* drives the system towards **II**_1_. The system quickly crosses **SP**, and stability is found within the **LC** and **Fix**_H_ segments.

Between **II**_2_ and **II**_3_ we find limit-cycle behavior (**LC**) that includes *P* = 0. Within the **LC** segment’s positive *P* domain, oscillatory power decreases with *P*. And **LC** segments terminate at 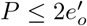, before **II**_3_ at *P* = 17 Hz. This ensures that large inputs 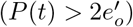 can drive the system out of limit-cycle behavior. If the input, *P*, is larger than Dom(max(**LC**)) < **II**_3_, then the system will converge to **Fix_H_**. However, for *P* ≥ 17 Hz, the system enters into the **II**_3_ segment and immediately shoots up to the **steady-state response segment**, 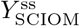, the higher **Fix**_H_ segment. For such inputs into the Jansen-Rit system, because there is no nonlinearity applied to inputs, no such saturation occurs.

### Energy Model of Underlying Dynamics

Although the bifurcation diagrams presented in Fig. 2 provide a qualitative overview of the dynamical features most relevant to the presence of intrinsic oscillatory behavior and its dissolution with external drive, they do not offer any insights into the internal dynamics fundamental to those features. Therefore, to address the underlying dynamics, we will introduce two new functions in our analysis: the system’s gross excitation function, *U*_sys_ (*Y*) (Eq. 62), and the excitation-inhibition work function, *ΣW*(*Y*) (Eq. 63). See Appendix E for the explicit derivation of *U*_sys_(*Y*) and *ΣW*(*Y*).

The gross excitation function, *U*_sys_(*Y*), determines the difference in the amount of “energy” stored in the excitation and inhibition firing rates of the network. Energy in this context is conceptually synonymous with the net excitation across the population under consideration. The population’s average firing-rate is therefore a reflection of the amount of this energy stored by the population. The excitation-inhibition work function, *ΣW*(*Y*), determines the amount of energy put into (or taken out of) the system at any point in the domain, *Y*. In this section, we will see how the interplay between *U*_sys_(*Y*) and *ΣW*(*Y*) determines the behavior of the system. For instance, given a particular **excitation landscape**, *U*_sys_, the proclivity for a transition to occur between two modes of behavior will depend on the sum of the excitation-inhibition work, *ΣW*, performed by the network near a transition point.

Before examining the standard Jansen-Rit system, and how the parameter modifications discussed in the previous section influence the system to satisfy the desired dynamics, we will define several terms that we will use in the ensuing analysis and discussion.

The system is in a **fixed excitation-state** when its output *Y^s^* = *V*_E_ − *V*_I_ is in an interval **Y_D_** such that 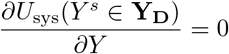. Therefore, fixed-excitation states are states such that small changes in *V*_E_ or *V*_I_ do not change the system’s net excitation, formally defined as for an *∊* > 0, the following holds: 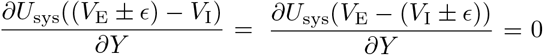. Fixed excitation-states occur because changes in the energy stored in the excitation firing rate of the network are offset by changes in the energy stored in the inhibition firing rate of the network, and vice-versa. Note that the segments **Fix_L_** and **Fix_H_** are each subsets of a respective interval of fixed excitation-inhibition states; that is: **Fix_L_** ⊆ **Y**d_L_ and **Fix_L_** ⊆ **Y**d_H_. Furthermore, fixed excitation-states are graded in terms of stability, and the most stable subset of fixed excitation-states occur when *U*_sys_(*Y^s^*) = 0, corresponding to a minimum in the system’s net excitation.

The system is driven internally by the “work” performed on it by the positive-feedback loop between the pyramidal and excitatory populations, and the negative-feedback loop between the pyramidal and inhibitory populations (see Appendix E for a more precise definition). The net work *ΣW* performed is therefore the sum of the work performed by these loops and the external drive. If *ΣW* ≠ 0, then a perturbation occurs in the system state, Δ*Y*. More precisely, if the system undergoes positive work then an increase in *Y* is generated, associated with an increase in *V*_E_, a decrease in V_I_, or both. **Stable equilibrium** occurs in a subset of fixed excitation-states, *Y*^*s*\*^, in which the work done on the system at *Y*^*s*\*^ is balanced, *ΣW* = 0.

### External Drive and Stable Oscillations

In the standard Jansen-Rit model with no external drive (*P* = 0), the excitation-inhibition work on the system becomes increasingly negative immediately to the right of the equilibrium point, 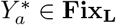 (Fig. 3A). Consequently, if the system starts at the equilibrium point, 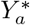, it will return to 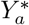 with small perturbations.

**Fig 3.**
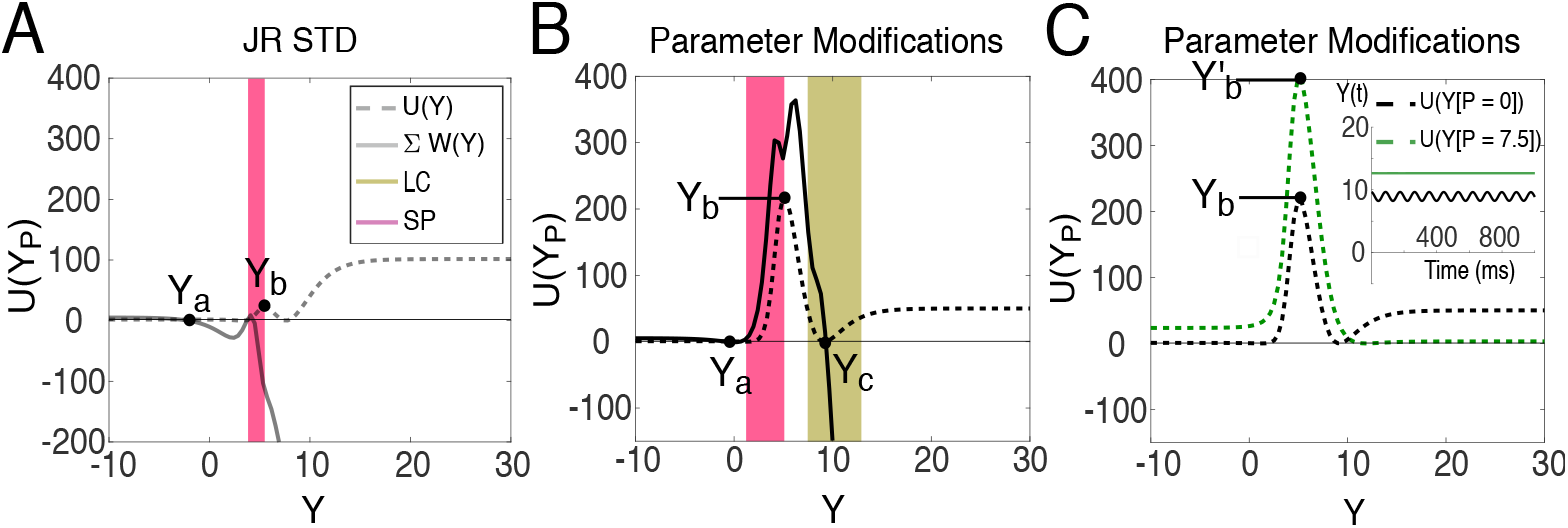
Interplay between the gross excitation function and the excitation-inhibition work function in the Jansen-Rit model. Excitation-inhibition work analysis over the standard Jansen-Rit model excitation-landscape in (A) and the parameter modified Jansen-Rit’s, 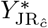, in (B), where *ĉ* = {120, 100, 20, 20}. The point *Y_a_* corresponds to the lower fixed point, **Fix_L_**, while the peak at *Y*_b_ corresponds to the separatrix region, and *Y*_c_ at the, *ΣW*(*Y*), zero-crossing corresponds to either the high fixed point, **Fix_H_** or the stable limit-cycle segment, **LC**. The standard Jansen-Rit system, under intrinsic drive, does not exhibit oscillatory behavior because the system’s excitation-inhibition work cannot overcome the excitation barrier (A). Therefore, neither **Fix_H_** or **LC** occur in the standard Jansen-Rit under this condition. On the other hand, the parameter modified system, 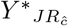, demonstrates an excitation landscape where sufficient net positive excitation-inhibition work is generated to overcome the energy-barrier, producing a **LC** within the energy-valley about *Y_c_*. In (C) the effects of external drive (*P* = 7.5Hz) on the excitation-landscape are shown for, *Y*^*^_*JRĉ*_. The energy-valley disappears and the system converges to **Fix_H_**.

By contrast, the unstable **SP** region is characterized by a peak in *U*_sys_, where 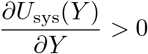, creating an **excitation barrier** at *Y_b_*, as indicated in Fig. 3A-C. To overcome this excitation-barrier, sufficient net positive work is needed to overcome it, *ΣW*(*Y* ∈ **SP**) *>U*_sys_(*Y* ∈ **SP**), but in the standard Jansen-Rit model, the excitation loop cannot produce sufficient net positive work and the inhibition-loop creates net negative work further preventing *Y* from increasing. Thus the only way to overcome the barrier is with strong external drive.

By increasing the excitatory coupling weights (*C*_EP_ and *C*_PE_) of the Jansen-Rit model, it is possible to strengthen the excitation-loop to produce sufficient positive-work on the system to overcome the energy-barrier in the **SP** region without large external drive. While increasing excitatory coupling also increases the excitation barrier (Fig 3B), the parameter regimes of the modified Jansen-Rit (Fig 2C) allows deviation from the fixed state.

Once over the **SP** energy-barrier the system begins to converge toward the equilibrium point 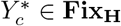, where *ΣW* undergoes a zero-crossing at the fixed excitation-state, 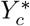. Overshoots or displacements, Δ*Y*, across an equilibrium point are hallmarks of higher-order dynamical systems (e.g. the Jansen-Rit system). Likewise, in order two or higher dynamical systems, these overshoots about a stable equilibrium point result in unstable oscillations. The work done on the system by the excitation-inhibition loops fluctuates between net-positive and net-negative work while 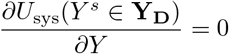, resulting in decaying oscillatory behavior that converges to 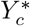 as *ΔY* → 0.

Stable oscillations emerge at a *ΣW* zero-crossing, *Y_c_*, within an excitation environment in which displacements about *Y_c_* in either direction produce 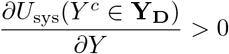. We refer to this convexity in the excitation-landscape as an **energy-valley**. Displacements, Δ*Y*, over *Y_c_* within this energy-valley such that Δ*U*_sys_ = *U*_sys_(*Y_c_* + Δ*Y*) − *U*_sys_(*Y_c_* − Δ*Y*) = 0, result in stable oscillations and the emergence of the **LC** segment. Ultimately, in the case of stable oscillatory behavior, the fluctuations in positive and negative work on the system are sustained by increases in the system’s net excitation, unlike in the case of decaying oscillatory behavior.

The effect of external drive on the excitation landscape, particularly the excitation-valley, is illustrated in Fig 3C. Two things occur simultaneously as external drive is added to the system. The first is the broadening of the excitation-valley, resulting in an increase in the displacement: Δ*Y*′ > Δ*Y* about *Y_c_*, such that Δ*U*_sys_ = *U*_sys_(*Y_c_* + Δ*Y*)′ − *U*_sys_(*Y_c_* − Δ*Y*′) = 0, within which stable oscillations occur corresponding to an increase in oscillatory power. The second is an increase in *Y_c_*, as indicated by the right-ward movement of 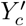, with the introduction of external drive in Fig 3C. Increases in *Y_c_* bring the system closer to the upper bound of *S*_F_. Eventually, the increase in oscillatory power is capped by the interaction between the oscillations and the upper bound of *S*_F_. Further increases in external drive continue to push *Y_c_* closer to the upper bound of *S*_F_ resulting in the reduction of oscillatory power.

The broadening of the excitation-valley with increasing external drive ultimately results in the elimination of the excitation-valley, and the reemergence of an interval **Y_D_** such that 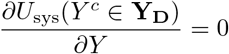. Accordingly, the zero-crossing point, *Y_c_*, becomes a stable equilibrium point once again, indicating the reappearance of **Fix_H_**. The system begins to exhibit decaying oscillations toward 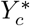, as before.

### E/I Balance and Stable Oscillations

In the bifurcation analysis we made several observations regarding the occurrence of intrinsic oscillations within a narrow range of excitatory-inhibitory coupling balance. In this section we will take a closer look at how oscillatory power varies over the 4-dimensional coupling coefficient space, **C_W_**, and use the energy approach to describe the geometry of the intrinsic oscillatory region in **C_W_**. In particular, we focus on identifying the constraints on network copuling necessary to produce strong intrinsic oscillations.

We first consider the 2-D cross-section of the full parameter space in which the inhibitory coupling weights vary (*C*_IP_ × *C*_PI_), and focus our attention on the subregion wherein intrinsic oscillatory behavior manifests, **C**_I*ϕ*_ (Fig. 4A). The geometry of **C**_I*ϕ*_ is determined by the bounded interaction between *C*_IP_ and *C*_PI_. We will delegate *C*_∑I_ to denote this interaction:

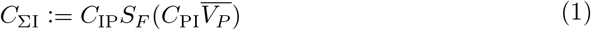

where 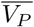 is the average pyramidal membrane potential. Intrinsic oscillations occur when *C*_∑I_(*C*_IP_, *C*_PI_) falls between a lower bound of 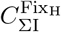 and an upper bound of 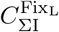:

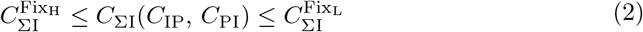

**Fig 4.**
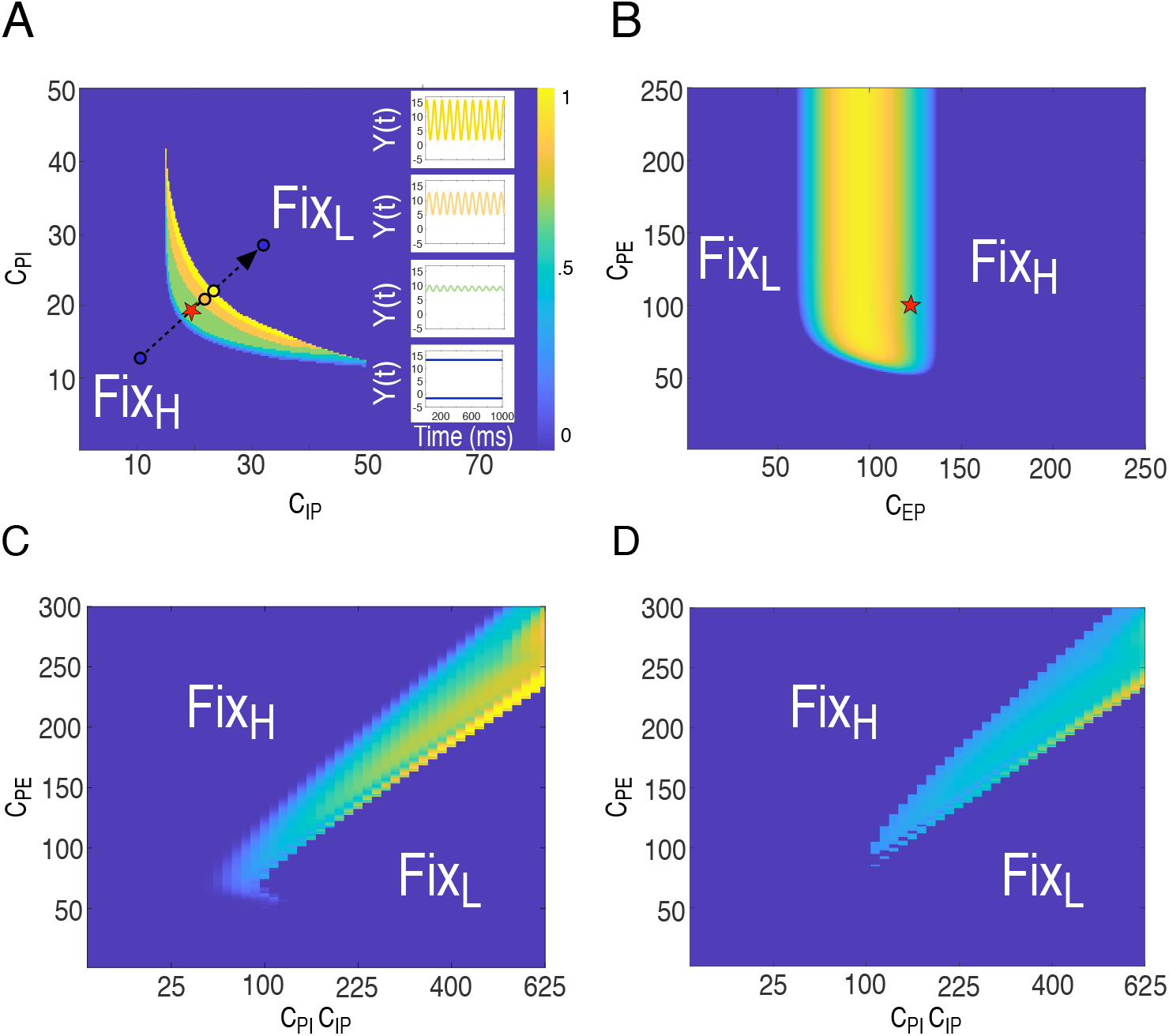
Dependence of alpha rhythms on E/I couplings. Normalized alpha-power (8-12 Hz) metrics over the inhibitory surface **C**_I*ϕ*_ (A) given 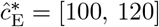 and **C**_E*ϕ*_ (B) given 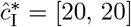. We map the coordinates of 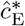 used to generate **C**_I*ϕ*_ in (B), and map the coordinates of 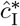 used to generate **C**_E*ϕ*_ in (A). In the inset of (A) we include simulations of four sampled points in **C**_I*ϕ*_. In (C) we generate the **C**_*E/I*_ map of the SCIOM system at intrinsic drive *P* = 0 with corresponding color map. Likewise in (D) we generate the **C**_E/I_ map of the SCIOM system at *P* = 5.

The upper 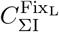 and lower 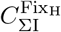 bounds within which oscillations occur were detected empirically in the bifurcation analysis according to the intersection of the **LC** segment with the vertical axis in the *Y* versus *P* phase-space. However, by fixing a point, 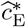, in the complementary surface, then the upper bound on the interaction between the two inhibitory coupling weights, 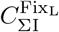, for intrinsic oscillatory behavior is determined as:

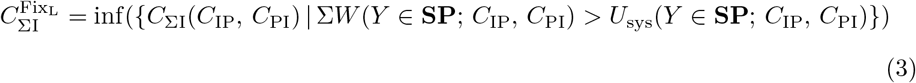

The excitatory-loop’s ability to produce sufficient net positive work to overcome the inhibitory-loop’s negative work and the excitation-barrier determines the upper bound on *C*_∑I_. If the coupling between the inhibitory and pyramidal populations exceeds 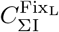, then intrinsic oscillations do not occur and the system will converge to **Fix_L_**.

On the other hand, the lower bound 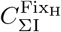 on *C*_ΣI_ for intrinsic oscillatory behavior is determined as:

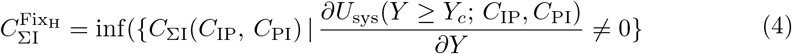

the smallest *C*_ΣI_ for which the excitation-landscape admits an energy-valley about *Y_c_*. The coupling between the inhibitory and pyramidal population must match or surpass 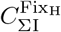 to support the presence of an excitation-valley in *U*_sys_(*Y*) about *Y_c_*. Otherwise, intrinsic oscillations do not occur and the system will converge to **Fix_H_**.

Alpha (8 to 12 Hz) power over **C**_IΦ_ indicates that increases in *C*_ΣI_ result in increases in the system’s oscillatory power, as shown in Fig. 4A. This observation is explained by the fact that: 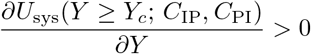 as 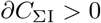; the rise in the excitation-valley results in an increase in the displacement, Δ*Y*′ > Δ*Y*, that the system undergoes about *Y_c_*, such that Δ*U*_sys_ = *U*_sys_(*Y*_c_ + Δ*Y*)′ − *U*_sys_(*Y_c_* − Δ*Y*′) = 0. Increases in *C*_ΣI_push the system to the left, along the *Y* coordinate in the energy landscape, *U*_sys_, driving the system away from the upper-bound of *S*_F_. Interactions with the upper-bound of *S*_F_ create the waxing and waning effect that is observed when increasing external drive or excitatory coupling as the system is pushed toward the upper-bound of *S*_F_. Consequently, because the system is pushed away from the upper-bound of *S*_F_ as inhibitory coupling is increased, waxing and waning of alpha power is not observed over **C**_IΦ_. Rather, the system’s oscillatory power is strengthened with continued increases in *C*_ΣI_ over **C**_IΦ_ until the system converges to **Fix_L_** before interacting with the lower bound of *S*_F_.

Next, we consider the 2-D cross section of the full parameter space in which the excitatory coupling weights vary (*C*_EP_ × *C*_PE_), with focus on the subregion wherein oscillatory behavior manifests, **C**_EΦ_ (Fig. 4B). Akin to **C**_IΦ_, the geometry of the excitatory surface, **C**_EΦ_, is determined by the bounded interaction between *C*_EP_ and *C*_PE_. We delegate *C_∑E_* to denote the interaction between the two excitatory coupling weights:

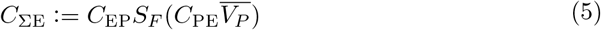

Intrinsic oscillations occur when *C*_EP_ falls between a lower bound of 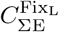 and an upper bound 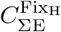:

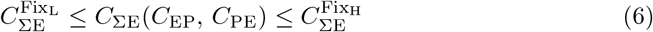

Fixing a point, 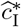, in the inhibitory surface, then the upper bound on the interaction between the two excitatory coupling weights, 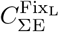, is determined as:

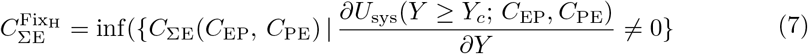

An excitation-valley is necessary for stable oscillations and increases in *C*_ΣE_ result in *U*_sys_(*Y* ≥ *Y_c_*) → 0, causing the system to converge to **Fix_H_**. Therefore, the annihilation of the energy-valley with increased coupling strength between the excitatory and pyramidal populations determines the upper bound, 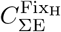, on 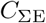.

On the other hand, the lower bound 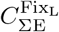 is determined as:

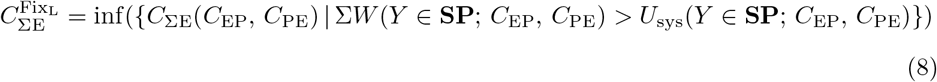

The excitatory-loop has to be strong enough to satisfy the excitation-barrier and to contend against the inhibitory-loop. This requirement imposes a lower bound 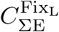 on *C*_ΣE_. Otherwise, the system will converge to **Fix_L_**.

Alpha power changes with **C**_EΦ_, for low *C*Σ_E_ alpha increases, and for high *C*_ΣE_ alpha power decreases, as shown in Fig. 4B. The dynamics behind this behavior is similar to the dynamics underlying the waxing and waning of alpha power that was observed when increasing external drive. Increases in *C*_ΣE_ result in the broadening of the energy-valley, which results in an increase in the displacement Δ*Y*′ > Δ*Y* that the system undergoes about *Y_c_*, such that Δ*U*_sys_ = *U*_sys_(*Y_c_* + Δ*Y*)′ − *U*_sys_(*Y_c_* − Δ*Y*′) = 0. Therefore, the oscillatory power of the system increase as *C*_ΣE_ is increased. At the same time, increases in *C*_ΣE_ push the system to the right, along the *Y* coordinate in the excitation-landscape, *U*_sys_(*Y*). Eventually, the increasing oscillatory amplitude begins interacting with the upper-bound of *S*_F_, imposing a cap on the height of the oscillations, and with the mean of the oscillations moving closer to the upper-bound of *S*_F_, the amplitude of the oscillations begins to shrink.

The similar dependence in **C**_IΦ_ on the two inhibitory coupling coefficients *C*_PI_ and *C*_IP_ results in the near symmetry seen in Fig. 4A. This occurs because of the mapping of 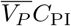 onto a near linear regime of *S*_F_, which matches the linear dependence on 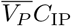. By contrast the dissimilarity in the dependence of **C**_EΦ_ on the two excitatory coupling coefficients *C*_pE_ and *C*_Ep_ results in the asymmetry seen in Fig. 4B. This occurs because 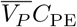, unlike 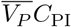, maps onto the nonlinear regime of *S*_F_.

The difference between the operating ranges of *C*_PE_ and *C*_PI_ arises from *A_i_τ_i_* ≫ *A_e_τ_e_*, and, as established earlier, the excitatory interaction must be strong enough to produce sufficient net positive work to overcome the excitation-barrier in order for the emergence of oscillatory behavior. Unlike C_PE_, C_PI_does not interact with the bounds of *S*_F_ within **C**_IΦ_. This is because *A_i_τ_e_* >> *A_e_τ_e_* mandates that large changes in *C*_PI_ toward either of the bounds of *S*_F_ drive the system out of the critical balance for oscillatory behavior defined earlier.

A representation of **C_W_** that allows us to evaluate the collective influence of the coupling coefficients on oscillatory power is attained by considering the ratio between *C*_∑I_ and *C*_∑e_:

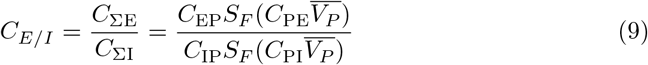

where 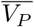 is the average pyramidal membrane potential. The space of *C_E/I_*, presented in Fig. 4C, substantiates the observations made in the bifurcation analysis regarding the occurrence of oscillations in a narrow range of excitatory-inhibitory coupling balance. The range of excitatory-inhibitory coupling balance associated with oscillatory behavior is bounded from above by 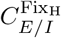, and from below by 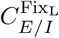:

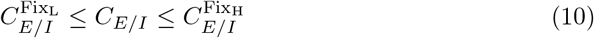

The upper-bound 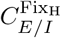 on *C_E/I_* for intrinsic oscillatory behavior is determined as:

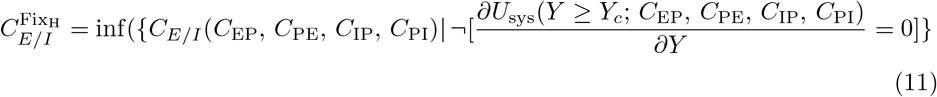

That is, a coupling balance 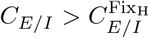 results in the system converging to **Fix_H_**; again, due to the annihilation of the excitation-valley. Conversely, the lower-bound 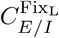 on *C_E/I_* for intrinsic oscillatory behavior is determined as:

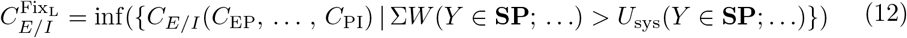

Where a coupling balance 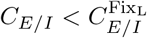 results in the system converging to **Fix_L_**; again, due to the inability of the excitatory-loop to satisfy the excitation-barrier.

Lastly, the effect of external drive on the oscillatory region in Fig. 4C is illustrated in Fig. 4D. External drive reduces the size of the oscillatory region, further narrowing the range of excitatory-inhibitory balance that generates oscillations. This is because increases in external drive cause *U*_sys_(*Y* ≥*Y_c_*) → 0, and cause the system to move closer to the upper-bound of *S*_F_.

### Oscillation Frequency

In the Jansen-Rit model’s formalism, the ratio between the average population capacitance, *A_η_*, over the lumped time constant, *τ_η_*, (incorporating the average membrane time constant, synaptic delays, and other temporal structures that shape the temporal dynamics) maps rates in Hz to synaptic drive in mV, 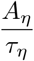. Together, the average population capacitance and the upper-bound on the firing-rate nonlinearity, determine the maximum synaptic drive, *V_F_*, to a population: 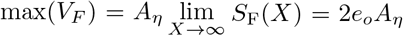. However, the average population capacitance and the lumped time constant are treated as independent quantities in the Jansen-Rit formalism. In the previous section, we saw that oscillations occur in a tight range of excitation-inhibition coupling balance, *C_E/I_*. We can extend this notion of a balance between excitation and inhibition underlying oscillatory behavior to synaptic drive readily:

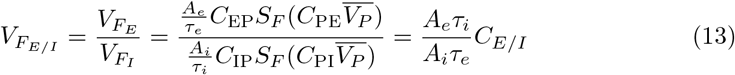

Likewise, as in the coupling coefficient discussion, oscillations occur within upper 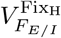 and lower 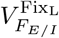 bounds:

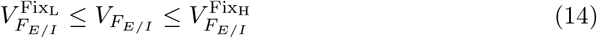

where 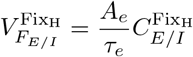 and 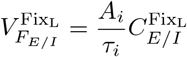, according to results from the previous section. In the Jansen-Rit’s treatment of *A_η_* and *τ_η_* as independent quantities, one can see how this balance can be easily disrupted. Consequently, when attempting to change the resonance structure of the system by changing the time constants, oscillations of different frequencies are not observed. The system either oscillates at 12 Hz or it does not oscillate at all across the majority of the parameter space. This is a critical limitation given the frequency variability of oscillations between different studies, and even different individuals. Likewise, the *A_e_* versus *A_i_* parameter space in Fig. 5A exhibits 10 to 12 Hz in a region that looks similar to the oscillatory region in Fig. 4C, but oscillations of different frequencies do not occur, and the system does not oscillate across the majority of the parameter space.

**Fig 5.**
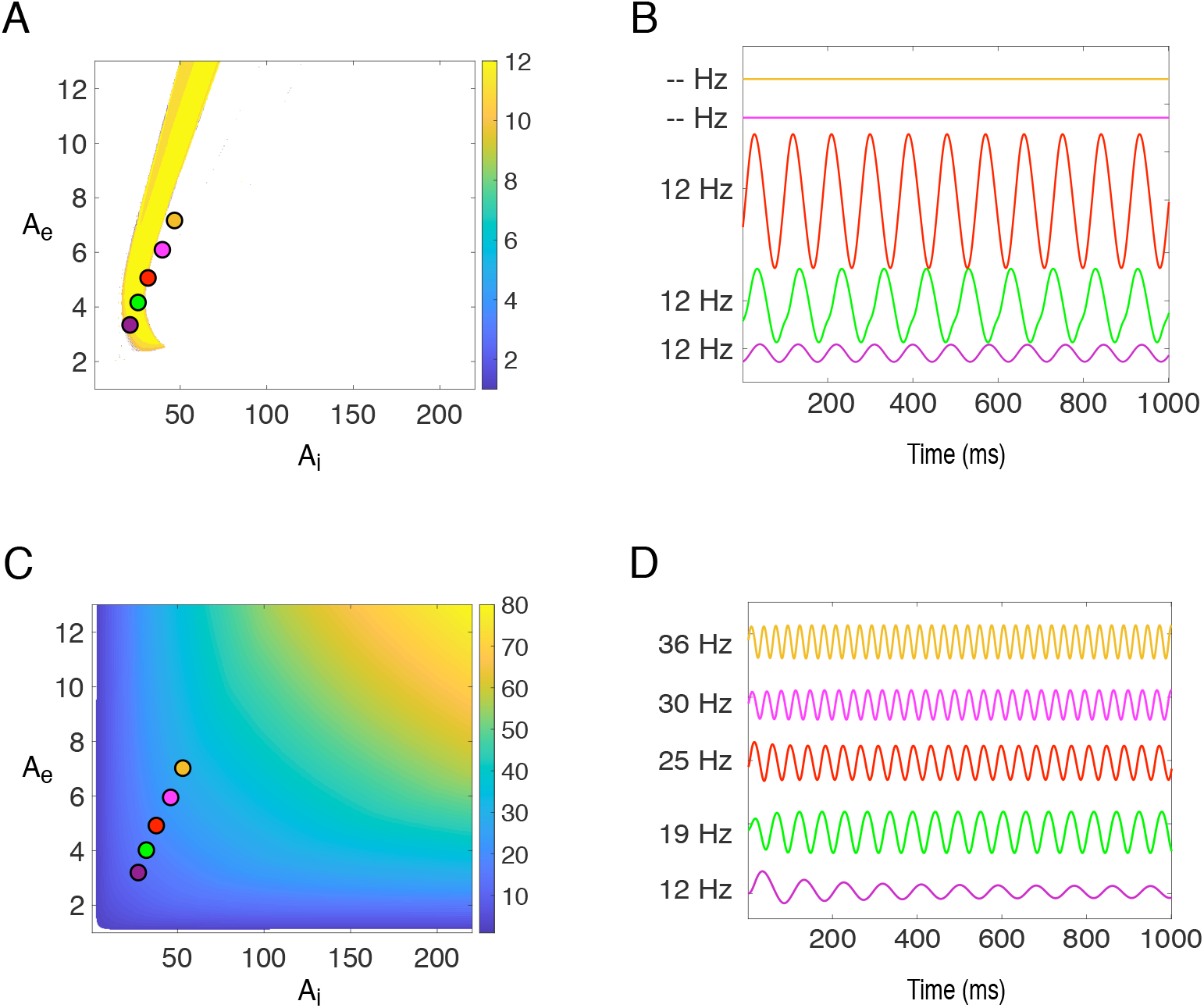
Dependence of oscillation frequency on E and I capacitances. The Jansen-Rit model’s frequency map as a function of excitatory and inhibitory capacitances (A). The SCIOM model’s frequency map as a function of excitatory and inhibitory capacitances (C). Sample points in the *A_e_* vs. *A_i_* capacitance space at coordinates: (3.25, 22), (3.75, 41), (4.29, 50.4), (5.00, 63.7), (5.89, 78.1), and (6.53, 92), were used in time-course simulations in (B) for the Jansen-Rit map, and in (D) for the SCIOM map.

In contrast, in the SCIOM model, the function 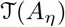 relates the average capacitance *A_η_* to the lumped time constant *τ_η_* of the population; therefore, the two quantities are dependent in this model. Their dependence, as defined in the methods section, is such that an increase in average capacitance results in an increase in the lumped time constant, and vice-versa. These two quantities are related by a conversion factor: 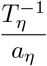 (see methods). Given two different capacitances, 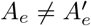, then the following holds true:

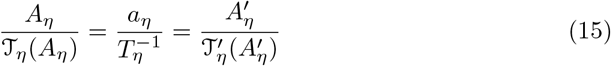

Therefore, for any choice of the conversion factor, 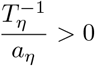, in which the balance between excitatory and inhibitory synaptic drive satisfies the bounds for intrinsic oscillatory behavior changes, in either population conductance or lumped time constants, will produce oscillatory behavior of a different frequency because:

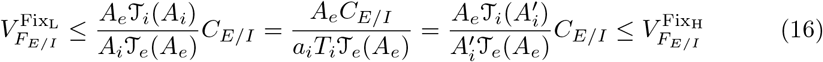

Consequently, by choosing a conversion factor such that the balance between excitation and inhibition drive satisfies Eq. 16 we can produce the frequency map shown in Fig. 5C by varying *A_η_* to generate oscillations of different frequencies but of similar spectral widths. Thus modest changes in the SCIOM parameters are sufficient to create different oscillation frequencies.

### Response Transients

A final advantage of the SCIOM model is the incorporation of synaptically mediated input drive. Given that inputs from a cortical microcircuit are mediated by synapses, this is of obvious benefit, but it also allows to accurately reproduce the non-linear response dynamics seen in physiological data. By contrast, even a parameter modified Jansen-Rit model that supports intrinsic oscillations (Fig 6B), is limited to linear responses. By contrast the SCIOM model (Fig 2C) responds to input with an appropriate non-linearity. The parameters are set such that input rate functions *P*(*t*) < 7 Hz are attenuated; hence, the system attenuates input function, *P*_1_(*t*).

**Fig 6.**
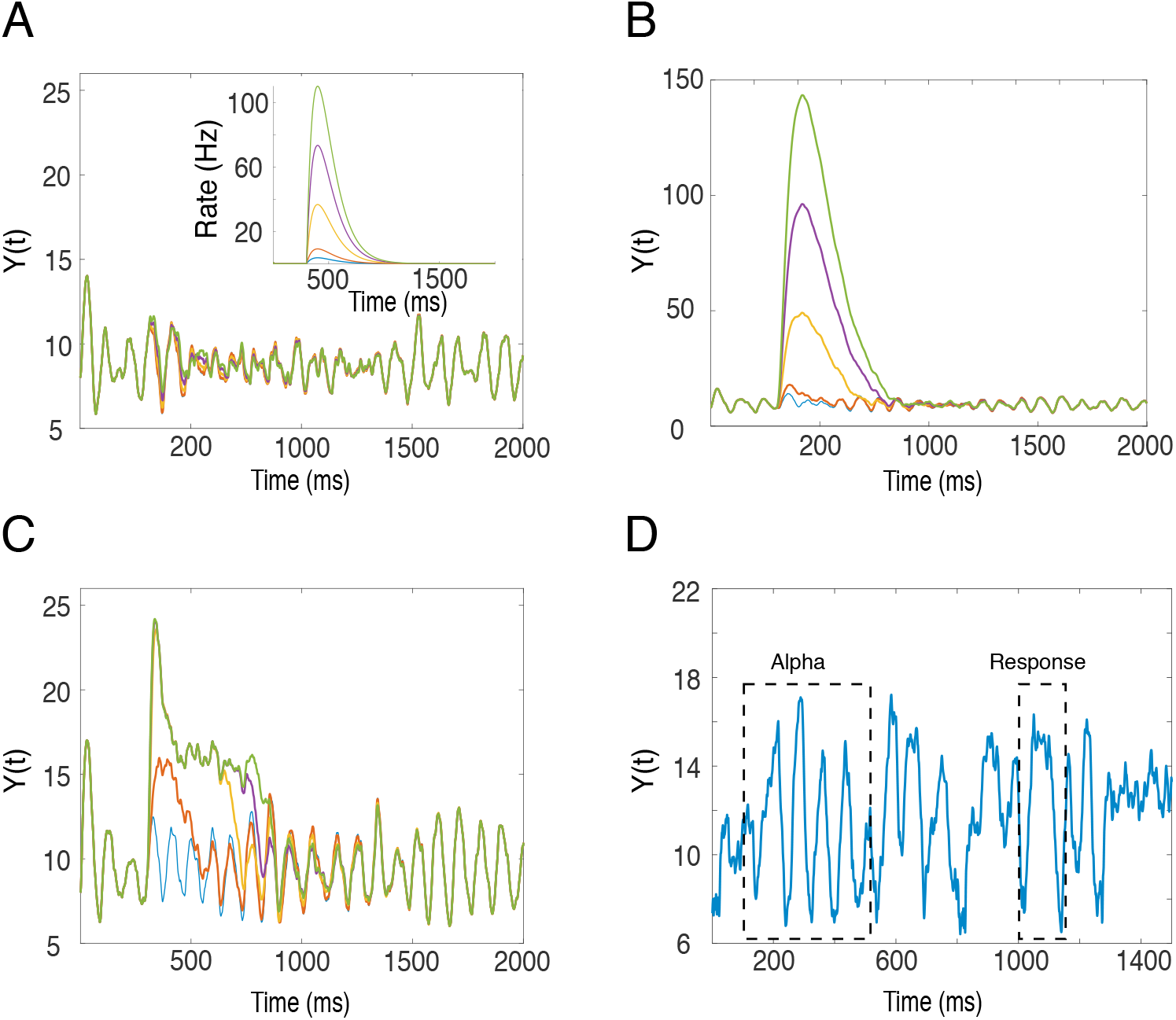
Transmission of input transients. The standard Jansen-Rit model’s response to the input functions presented in the inset of (A). The parameter optimized Jansen-Rit model, allowing the transmission of transients, response to the same input functions in the inset of (A) is presented in (B). The SCIOM model’s response to the same input functions is presented in (C). Raw LFP data over the V4 of a trained adult monkey performing a visual shape detection task illustrates prominent alpha-rhythms preceding the onset of a noisy stimulus in its visual space, followed by a shape stimulus, followed by V4 response, and a saccade.

The SCIOM model exhibits several modes of response depending on the state of the system and the strength of the input function. First, **subliminal responses** are those that are attenuated by the system. **Intermediary responses** are those that produce a sharp transient that are sometimes followed by a broader, decaying transient.

**Suprathreshold** responses ares ones in which the system generally produces an initial sharp transient that is followed by a steady-state response, 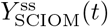.

Fig 6C illustrates the SCIOM model’s response to the set of five input-rate functions that were used in the preceding Jansen-Rit simulations in Fig 4. In this case, the model’s parameters were set to attenuate input-rate functions *P*(*t*) < 7. What we see is the model’s attenuation of *P*_1_(*t*), but to *P*_2_(*t*) it produces an intermediary response. This intermediary response consists of a sharp transient that’s followed by a slower decay. To input functions: *P*_3_(*t*), *P*_4_(*t*), and *P*_5_(*t*), the model responds with a suprathreshold response consisting of an initial sharp transient, that is followed by steady-state components. The steady-state component for *P*_5_(*t*) lasts the longest. These response dynamics are compared to a segment of a raw V4 LFP recording taken from a behaving adult male Macaque, performing a visual task (Fig 6D). A segment of strong alpha oscillations precedes the onset of a visually evoked transient that consists of a sharp transient followed by a steady-state response, comparable to the SCIOM model’s suprathreshold response behavior.

### Interaction between intrinsic oscillations and transient responses

In this section, we will examine the SCIOM model’s steady-state response component as a function of the internal state of the system (its alpha-power), and examine the initial sharp transient component as a function of alpha-phase. From here on, for brevity, we will use the term **fast-component** when referring to the initial sharp transient of the response. Furthermore, we will use the term **oscillatory state** in reference to the power of alpha in the system.

We can solve for the SCIOM’s steady-state response function, 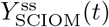, precisely by taking the upper limit of *S*_F_ in Eq. 34:

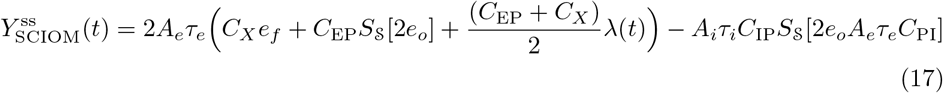

By examining Eq. 17, we see that the steady-state behavior, 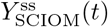, is strongly determined by the inhibitory influence. This is due to the fact that *A_i_τ_i_* ≫ *A_e_τ_e_*.

Fig. 7A illustrates the SCIOM model’s responses to a visual input in the presence of 3 phase aligned oscillations of different strengths (weak (Ψ_W_), moderate (Ψ_M_), and strong (Ψ_S_)). The three oscillations were obtained through changing coupling interactions in the network. The respective steady-state (post transient) responses 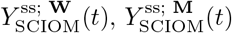, and 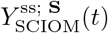 are anticorrelated with the oscillatory state of the system: the greater the alpha power of the system, the smaller the amplitude of its steady-state response, and vice-versa.

**Fig 7.**
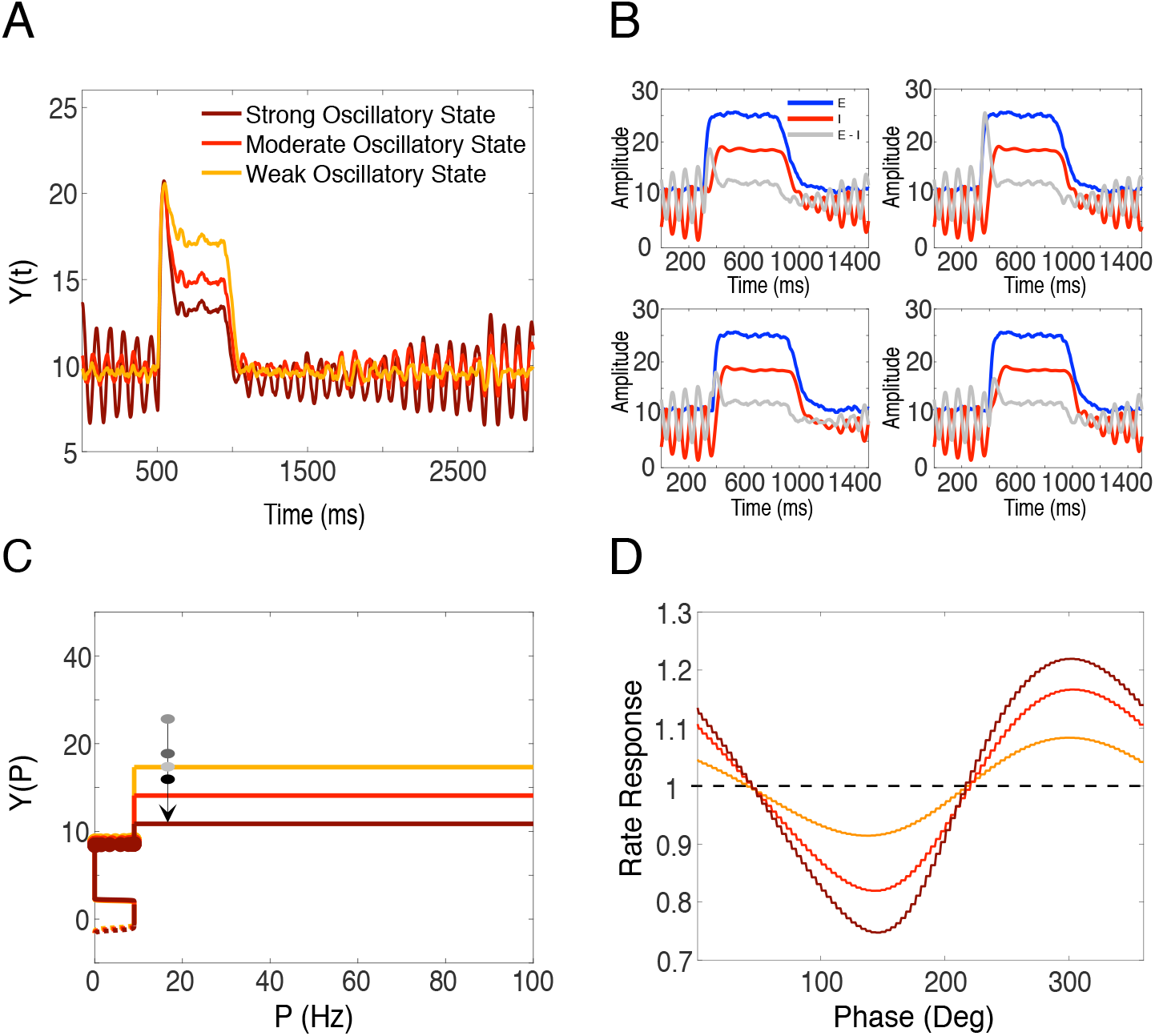
Alpha phase dependence of transient responses. The SCIOM model’s response to a fixed input function at three different oscillatory states: strong, moderate, and weak, with all other parameter conditions equal, induces three different steady-state response amplitudes (A). Steady-state response amplitude is also anti-correlated to the power of alpha oscillations in the system. In (B) the SCIOM model is presented with the same input-function in (A) but at four different phases: 0, 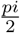, *π*, and 2*π*. The phase effect is reflected in the amplitude of the initial transient that precedes the steady-state response, and is related to the time lag in the inhibitory population’s response in curving excitation. The phase-space representation reconciles both the phase-dependent and the amplitude dependent effects (C). The initial transient represents an overshoot across the equilibrium segment 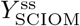, and the height of the overshoot is phase dependent. The system then eventually settles back down to equilibrium in 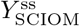. In (D) we demonstrate the full phase response curve associated with the three oscillatory conditions in (A). The phase-dependency scales with oscillatory power.

To explain these observations, we take another look at the *Y* vs. *P* phase space (Fig. 7C). The mean of the steady-state function, 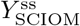, corresponds to a segment of equilibrium points in excitation-inhibition, E - I. The trajectory of the system in its phase space when undergoing a suprathreshold response involves an initial displacement from excitation-inhibition equilibrium at some *P* ≥ **II**_3_ corresponding to the **fast-component** of the response. When the system’s phase space trajectory falls back down to the equilibrium curve, if *P* ≥ **II**_3_ still holds, then the system maintains a steady-state response, 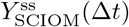 for the duration that *P*(Δ*t*) ≥ **II**_3_. Moreover, 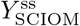 is negatively correlated with alpha-power simply because inhibition, as was established earlier, is positively correlated with alpha-power, and increased inhibition 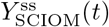.

The response of the SCIOM model to the fast-component is dependent on the phase of the alpha rhythm, as shown in Fig. 7B. In this figure, we also look at the excitatory and inhibitory time courses, and show that the excitatory population’s response to *P*_4_(*t*) at different phases is virtually identical. By contrast, the inhibitory population displays a phase dependent time lag in generating a response. This results in a phase dependent population response (Fig. 7D), as defined by the max amplitude of the fast-component at a particular phase over its average maximum amplitude across all phases. The phase response of the system is also dependent on the power of the oscillations. A larger phase effect is seen for the strong oscillatory state than for the moderate and weak, in that order.

## Discussion

In order to understand how the prominent low frequency 8 to 12 Hz rhythms in cortex impact information processing, we adapted a classic neural mass model (Jansen-Rit) to model a cortical microcircuit which produces realistic activity dynamics and relies upon physiologically derived parameters. Our motivation was to address two critical issues in the original model: the absence of low frequency oscillations without strong input drive and the inconsistency between input drive and output activity. Both of these inconsistencies were addressed with modest changes in connectivity parameters and the addition of a input non-linearity consistent with the synaptically mediated inputs that cortical neurons receive and that govern connections within the microcircuit. We further demonstrate that these modest changes produce realistic non-linear responses to input transients and that, consistent with experimental observations, the magnitude of those responses depends on the phase of alpha rhythms. By deriving a potential energy approach to understanding oscillatory dynamics, in which stable oscillations occur in an energy valley, we found that in our model, unlike the original Jansen-Rit model, increasing external drive results in the reduction and ultimate disappearance of this potential energy valley, thus allowing strong input transients to be faithfully transmitted. These results are robust to a range of intracortical connectivities, but are critically dependent on the ratio of excitation to inhibition.

The presence of strong oscillations in the absence of strong input is consistent with the intrinsic oscillations reported in cortical slice preparations as well the prominence of alpha oscillations in states of relatively quiescence. Indeed, the first reports of such oscillations by Berger in 1930s [6] noted that such oscillations in visual cortex were most prominent when eyes were closed and largely absent when subjects were visually engaged. Decades of subsequent study have demonstrated that the oscillations are associated with a lack of attention and a corresponding drop in visual performance [39, 64, 52, 22].

In our model, oscillations are intrinsic to the circuit [20], and not reliant on an external oscillatory signal, consistent with multi-site recordings in V4 and FEF [7]. However, critical features of the cortical microcircuit, including laminar heterogeneities in both connectivity and alpha rhythms [62], which are thought to originate infragranularly and synchronize across laminae, are absent in the Jansen-Rit model.

While this circuit has distinct populations of projection pyramidal cells and excitatory and inhibitory interneurons, it is clearly a simplification of the rich heterogeneity of cell types and neurotransmitters in the cortex. While structural heterogeneity can strongly affect network dynamics [37], recent evidence suggests that excitation/inhibitory ratios in pyramidal cells are conserved at a cellular level [68], which is consistent with the critical dependence of our model on these ratios. Excitatory/inhibitory imbalance has been associated with cortical dysfunction both experimentally [16, 19, 33, 26] and theoretically [9, 42, 70, 3, 49, 48, 69, 58, 12, 54].

The SCIOM model is robust to modest variations in the effective decay constant of excitatory and inhibitory postsynaptic potentials. It is also robust to large variations in inhibitory and excitatory coupling as long as their ratios are maintained. This is consistent with recent studies in invertebrate model systems suggesting that it is network behavior [50, 11], rather than the exact mixture of connections and neurotransmitters, that is strongly conserved across systems and individuals. In our model, significant excitatory/inhibitory imbalance leads to fixed points in the which the system is “stuck” and unable to faithfully transform input transients. This is consistent with the premise that excitatory/inhibitory balance is strongly conserved and necessary for the accurate processing of incoming information in both sensory and cognitive contexts.

Our model critically depends on input non-linearity. Non-linear input-output relations, in addition to being physiologically well motivated given the synaptic nature of inputs to a cortical microcolumn, have long been associated with critical functioning including the maintenance of onset transients throughout a heirarchical system and allowing for stochastic resonance. By contrast, the standard Jansen-Rit system does not transmit inputs at all as they get attenuated by the ongoing oscillations and even the parameter modified Jansen-Rit system responds linearly to input transients. This is particularly problematic when considering the high input case in which, in the absence of any output thresholds, output increases without limit.

In addition, SCIOM’s response to input transients is strongly dependent on the phase of alpha oscillations [60, 10], consistent with a variety of psychophysical studies involving visual, auditory, and somatosensory detection [40, 38, 44, 18] and discrimination [25, 55, 5], and physiological studies highlighting the role of ongoing activity in explaining response variability [34, 29]. The physiological basis for these effects has long been the subject of modeling [51, 40, 41]. For example, models have typically assumed that the inverse relationship between alpha waves and cortical excitability is governed by a strong association between alpha waves and inhibitory neurons [36, 43], and that control of alpha wave magnitude is governed by thalamocortical inputs [63]. However, a variety of studies have pointed to an association of top-down feedback inputs [56, 46, 4], which largely target excitatory cells [24], as critical to the regulation of intrinsically generated alpha power [8].

SCIOM is consistent with such observations: the interactions between alpha and input transients are a network phenomena, and are not solely due to a selective action upon the inhibitory population. Changes in response transients occur because of a difference of response latencies in the excitatory and inhibitory populations.

SCIOM makes an interesting prediction regarding the effects on alpha upon the response dynamics to brief input transients: the sustained response that follows the initial transient is dependent on the oscillatory state of the system, but, unlike the initial response, not its phase. This is because the oscillatory state of a system is a reflection of the underlying excitation-inhibition balance, and the amplitude of the steady-state response is a function of the that balance. While such effects on response dynamics are certainly consistent with experimental observations pointing to a complexity of response effects with ongoing alpha rhythms [35, 28], verification of this prediction would depend on delivery of impulse-like changes in input rates, which is unlikely to be observed, even in the case of brief external inputs, in the case of sensory transduction mediated stimulation, and may be require direct phase-specific electrical stimulation.

Because a major motivation of our approach was to impose consistency between input and output regimes, a natural extension of would be construct 2-D arrays of these microcircuits to study the spatiotemporal interactions between alpha and stimulation across the surface of the cortex and thus make predictions about the spatiotemporal generation and propagation of macroscopic signals that give rise to EEG observations. Finally, should these studies result in data consistent with existing experimental data, the model would have considerable predictive value for the design of multi-dimensional microstimulation paradigms.

## Methods

### Computational Models

This section begins with a description of the Jansen-Rit model as originally defined, placing particular emphasis on the two transformations that the model performs. Then, we will introduce a third transformation that leads us to the SCIOM model whose dynamics better simulates realistic response transients and alpha rhythms seen in experimental data. Afterwards, we will order reduce the SCIOM model to arrive at a single expression that encapsulates the entire equilibria set of the model.

### Jansen-Rit

The Jansen-Rit (JR) system models a local cortical population with a set of three, coupled, nonlinear ordinary differential equations of the second order that describe the mean dynamics between three interconnected neuronal populations, namely: excitatory (E), inhibitory (I) interneurons, and pyramidal (P) long range projection neurons.

In essence, the model approximates the dynamics of the aforementioned populations as an equality between the second order derivative of their average membrane potentials and their received net synaptic drive. In this process, each population performs two transformations. The first is a transformation from membrane potential to **firing-rate** and the second is a transformation from pre-synaptic action potential rate **input-rate** to postsynaptic potential.

A block diagram of the connectivity between these populations is shown in 1A. While the mathematical definition of the model is as follows. Given *t* ∈[0; ∞):

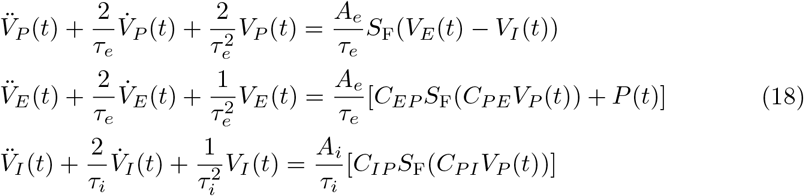

The left hand sides of the equations determine the membrane potential dynamics of the neuronal populations. The right hand sides describes their interactivity with each other and with external sources, in terms of firing-rates. The average voltage (mV) for each population is *V_j_* (*t*) for *j* ∈ {*P, E, I*}and its first and second derivatives, 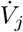 and 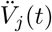. The lumped parameter *τ_η_* for *η* ∈ {*e, i*} combines both the membrane time constant and the synaptic delay together, for the excitatory (e) and inhibitory (i) populations, respectively. Aηscales the net input-rate and determines the maximum synaptic response to an impulse of input (see, “Transformation 2,” of the Computational Models section). The ratio 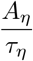 converts the net input-rate to an electric potential, which in the Jansen-Rit model is unbounded. Lastly, the parameter *C_γ_* is the coupling coefficient between the different populations where *γ* ∈ {PE, EP, IP, PI}, and PE represents the drive from the pyramidal population to the excitatory population. In the Jansen-Rit model, these parameters: *C*_EP_, *C*_PE_, *C*_IP_, and *C*_PI_ are each proportional to a global coupling parameter *C_G_*. That is *C_γ_*: = *ρ_γ_*, where *ρ_γ_* is a scalar between 0 and 1, see 1.

#### Transformation 1: Membrane Potential to Firing-Rate

The input drive to each population from the other populations is transformed through a sigmoid function *S*_F_ from a potential to an average firing-rate in Hz. We’ll refer to *S*_F_ as the **firing-rate nonlinearity**:

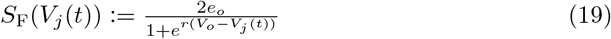

where *e_o_* represents the half maximum firing-rate of the population (Hz), *V_o_* is the potential (mV), that generates the half maximum firing-rate, and *r* describes the average membrane sensitivity (mV^−1^). The coefficient 2*e*_0_ determines the upper-bound of the sigmoid output rate, and *r* is the slope at the half maximum point, *V_o_*.

The external input to the system is *P*(*t*), representing “pulse-density” (Hz), carries both upstream information and noise in the Jansen-Rit formulation. We will separate the input-rate function from intra-network sources, *F_N_*(*t*) := *S_F_*(*C_γ_V_j_*(*t*)), and extra-network sources, *F_X_*(*t*) : = *P*(*t*). Hence, the net input-rate function to a population is the superposition of the intra-network and extra-network input-rate functions:

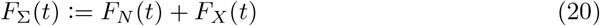

All the standard values of the Jansen-Rit model’s parameters are defined in 1. These parameter values were chosen by the authors of the Jansen-Rit to generate stable alpha-rhythms.

**Table 1.** Standard values of the Jansen-Rit model’s parameters.

| PSP | | CC's | | $S_F$ | |
| --- | --- | --- | --- | --- | --- |
| Param. | Value | Param. | Value | Param. | Value |
| $A_e$ | 3.25 mV | $C_{EP}$ | 125 | $e_o$ | 2.5 Hz |
| $A_i$ | 22 mV | $C_{PE}$ | 100 | $V_o$ | 6 mV |
| $\tau_e$ | 10 ms | $C_{IP}$ | 20 | $r$ | $0.56 \text{ mV}^{-1}$ |
| $\tau_i$ | 10 ms | $C_{PI}$ | 20 | | |

#### Transformation 2: Input-Rate to Membrane Potential

The model’s synaptic impulse response *H_η_*(*t*), (see Appendix A for derivation) is:

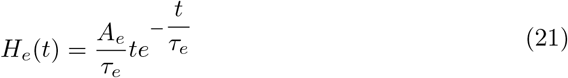

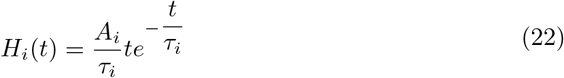

for the excitatory and inhibitory populations respectively. The excitatory postsynaptic-potential (EPSP) and inhibitory postsynaptic-potential (IPSP) impulse responses are shown in Figure 8.

**Fig 8.**
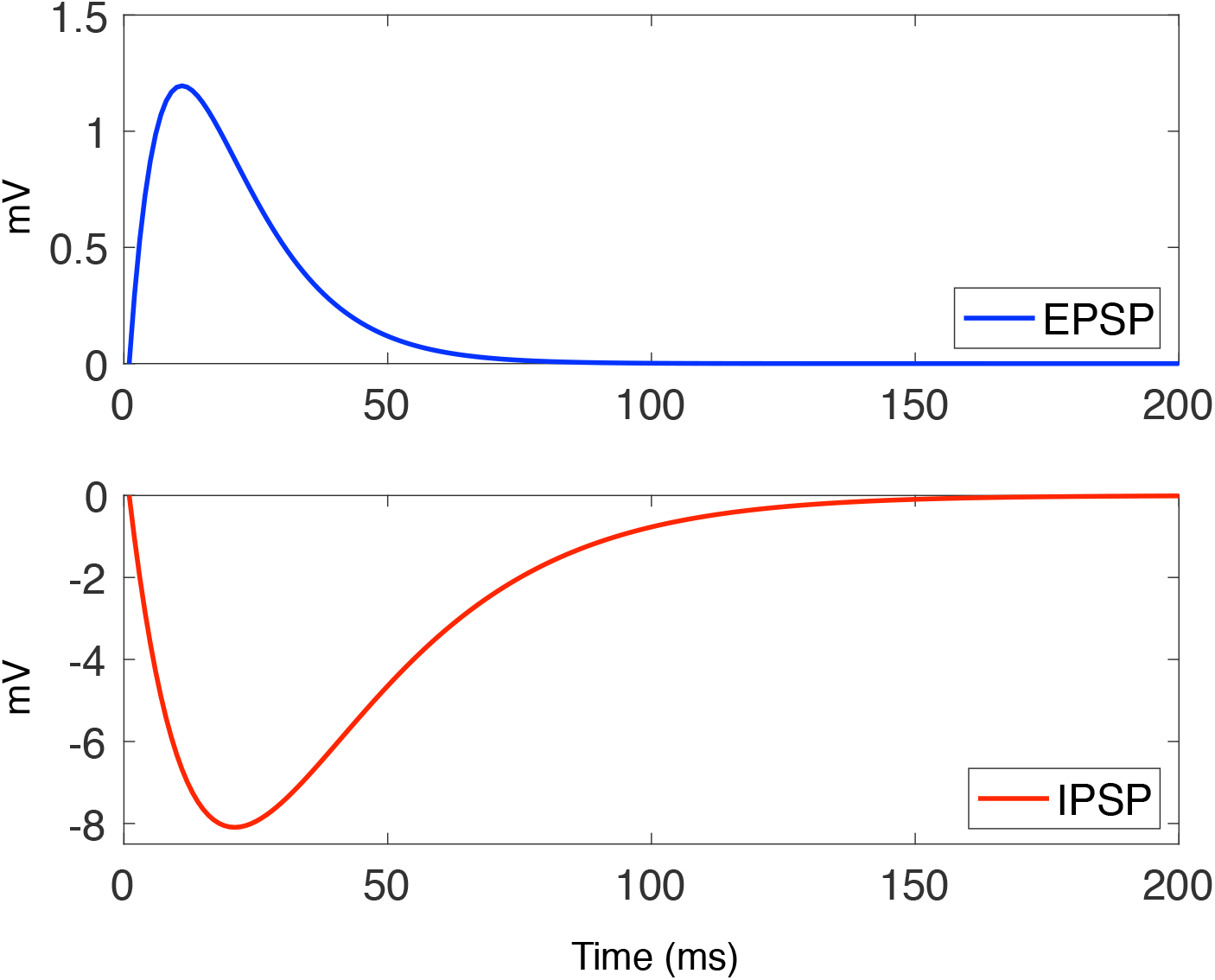
Impulse response of the Jansen-Rit populations’ EPSP (blue) and IPSP (red). The coefficients *A_e_* = 3.25 mV and *A_i_* = 22 mV scales the max response. The time constants are *τ_e_* = 10 ms and *τ_i_* = 20 ms.

Thus, from functional analysis, the membrane potential of a population *j* at time *t* is the convolution of its impulse response function *H_η_*(*t*) with the net input-rate function *F*_Σ_(*t*) to the population:

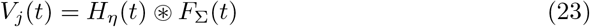

Lastly, the output *Y* of the model approximates the measured local field potential of the microcircuit as the sum of the excitatory and inhibitory population potentials:

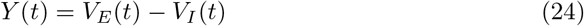

### Equilibrium Expression for 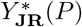

An equilibrium equation for the Jansen-Rit model relating critical states 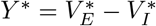 to external input-rate *P* has been previously presented [23, 2]:

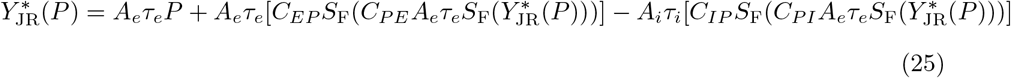

Equation (25) defines a relation from *P* to *Y*\*. The derivation of this expression involves standard order-reduction of the system (18) and substitution. We will not show these steps for the Jansen-Rit system; however, we will explicitly derive an equilibrium equation for the adapted model in the ensuing sections.

### SCIOM (Self-consistent intrinsically oscillating microcircuit) model

The SCIOM model differs from the Janset-Rit model in 3 aspects (1): its input drive is synaptically mediated, there is a synaptic capacitance, and several coupling parameters are changed.

#### Synaptically mediated input

The first change from the Jansen-Rit model is how net-input rates *F*_Σ_(*t*) are treated and their linear transformation to synaptic drive.

Let *V_F_*(*t*) denote the synaptic drive. In the Jansen-Rit model, the conversion factor 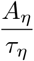 maps rates in Hz to synaptic drive in mV linearly:

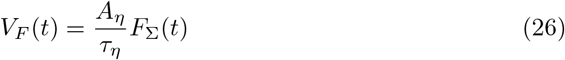

Furthermore, because the membrane-potential dynamics of population *j* responds linearly to synaptic drive:

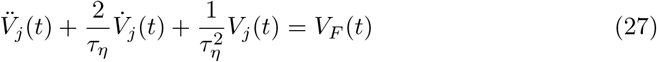

it therefore responds linearly to net input-rates *F*_Σ_(*t*) as well. There are three issues with this behavior. First, it allows the populations to respond to physiologically impossible negative input-rates. Second, there are no physiological constraints or bounds on the synaptic drive and extreme events are possible. Although one could avoid these predicaments by constraining *F*_Σ_(*t*) within a physiologically plausible bound the populations will still respond linearly to *F*_Σ_(*t*), and the output of the model becomes a scaled version of the input.

This **response linearity** is neurophysiologically inconsistent with the nonlinear nature of synaptic mediated dynamics. **Response nonlinearities** are necessary for multiple stable states and the ability to transition to distinct modes of response dynamics. Synaptic nonlinearities can also attenuate subthreshold inputs, such as noise, to enhance relevant features resulting in improved noise rejection and information processing.

To address the issues caused by the linear responses in the Jansen-Rit model, we will add a nonlinear transformation function 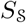 to the synaptic drive, that maps presynaptic firing-rates in Hz to the rate of synaptic events in Hz. We’ll refer to 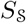 as the **synaptic nonlinearity**:

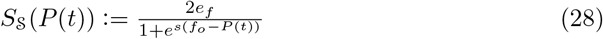

here *e_f_* represents half maximum input-rate (Hz), *f_o_* is the input-rate (Hz) that achieves half the maximum synaptic drive, and *s* is the synaptic membrane sensitivity (Hz^−1^) to input-rates. The coefficient 2*e_j_* determines the upper-bound of 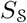, and *s* corresponds to the slope at the half maximum point, *f_o_*.

The nonlinear transformation 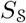 on the input-rate functions *F_N_*(*t*) and *F_X_*(*t*) leads us to a new model of synaptic drive to population *j*:

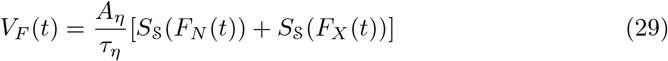

Consequently, the membrane-potential dynamics of population *j*, which depends linearly on their synaptic drive *V_F_*(*t*), as defined in equation (27), depends nonlinearly on input-rates. This change leads us to the “Self-consistent intrinsically oscillating microcircuit” (SCIOM) model. The block diagram of this new model is provided in 1B. The corresponding mathematical definition of the SCIOM model is as follows:

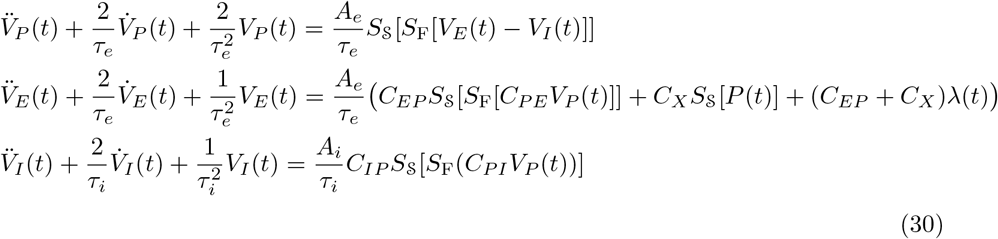

where λ(*t*) is a Poisson distributed random variable simulating exogenous noise. The parameter *C_X_* weighs the average synaptic drive from external interactions. Observe that increasing either *C_EP_* or *C_X_*, the strength of the feed-forward process, increases the system’s noisiness.

#### Capacitance

The next significant change from the Jansen-Rit model concerns the relationship between the lumped time constant *τ_η_* and the synaptic capacitance *A_η_* of a population. The Jansen Rit model treats them as independent quantities: changes in *A_η_* do not result in changes in *τ_η_*. By contrast, the SCIOM model treats these two quantities as dependent in which the time constant of a neuron should be approximately proportional to its synaptic capacitance, as a rough estimation:

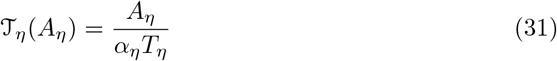

Here *α_η_T_η_* serve as a conversion factor that converts a given synaptic capacitance *A_η_* to a particular time constant 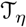. In this paradigm, changes in the average synaptic capacitance of a population result in a proportional change in its lumped time constant. This is physiologically realistic because, on average, as the capacitance across a population of neurons increases the amount of voltage change across the population increases on average, and the length of time it takes for its restoration to baseline increases as a result. Consolidating equation (31) with system (30):

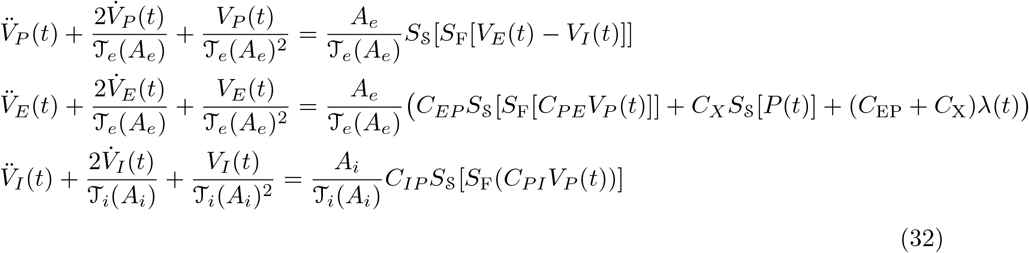

Different oscillations of similar spectral-width but of different frequencies can be generated through different choices of synaptic capacitance, *A_η_*. Alternatively, with this formulation, oscillations of different spectral-width but of the same frequency can be produced by varying the conversion factor *α_η_T_η_*, while holding the synaptic capacitance, *A_η_*, fixed.

#### SCIOM parameter values

For convenience, all parameter modifications of the SCIOM model in this manuscript will be in reference to 2. When presenting a parameter modified SCIOM model, we will list the parameter changes that are different from the list of parameters presented in this table.

**Table 2.** Values of the SCIOM model’s parameters.

| PSP | | CC's | | $\mathbf{S}_F$ | | $\mathbf{S}_S$ | |
| --- | --- | --- | --- | --- | --- | --- | --- |
| Param. | Value | Param. | Value | Param. | Value | Param. | Value |
| $A_e$ | 3.25 mV | $C_{EP}$ | 125 | $e_o$ | 2.5 Hz | $e_o$ | 2.5 Hz |
| $A_i$ | 22 mV | $C_{PE}$ | 100 | $V_o$ | 6 mV | $V_o$ | 6 mV |
| $\tau_e$ | 10 ms | $C_{IP}$ | 20 | $r$ | $0.56 \text{ mV}^{-1}$ | $r$ | $0.56 \text{ mV}^{-1}$ |
| $\tau_e$ | 10 ms | $C_{PI}$ | 20 | | | | |
| | | $C_X$ | 40 | | | | |

### SCIOM’s Equilibria Set

Because of the nonlinearieties in the SCIOM model, an analytical solution to Eq. 30 is unattainable. Nevertheless, analysis of the equilibria set can reveal behaviors exhibited by the model. Here we derive a minimal set of expressions that reduces the model’s six-dimensional state-space that relates equilibria *Y*\* to the external constant input-rate function *P*.

### Equilibrium Expression for 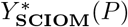

We begin with the reduced-order representation of the SCIOM model, where the higher-order terms are expressed as first-order expressions of time, *t*:

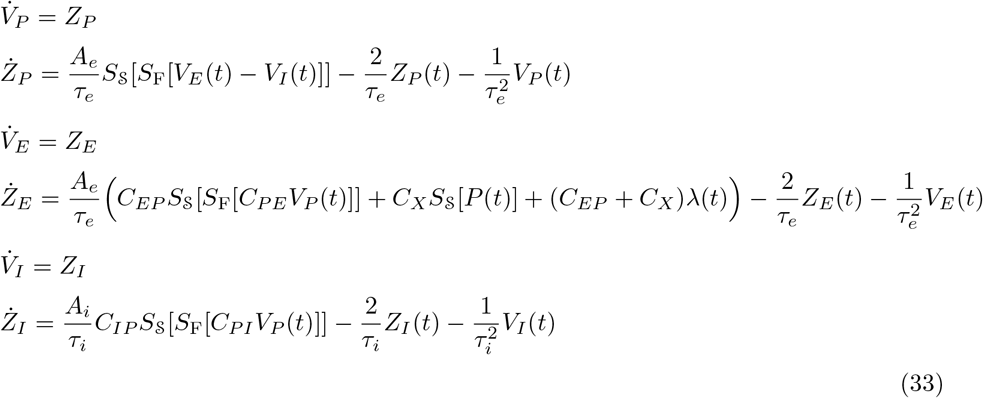

Equilibrium occurs within a subset (limit-set) of the solution space of (33) when the differential terms become 0. We will refer to this space as the space of **critical-states**. We can solve for said solution space, but uncovering the limit-set requires the use of a bifurcation analysis software (we will use XPP-AUT). Nevertheless, solving for critical states: 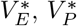, and 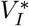, results in the system:

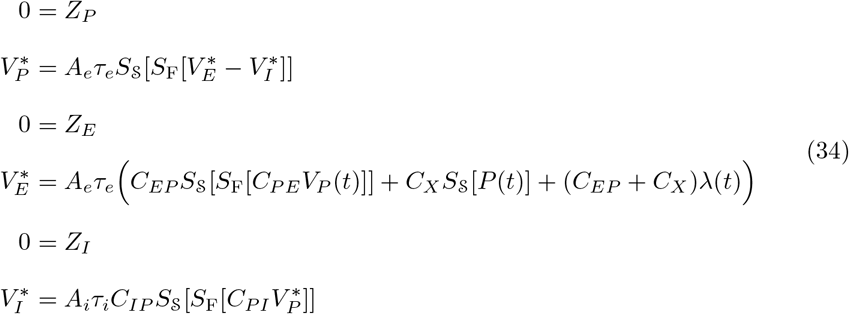

The equilibria set defined by (34) is 6-dimensional, however, three of these dimensions are trivial: *Z_P_* ≡ 0, *Z_E_* ≡ 0, and *Z_I_* ≡ 0. Thus, proceeding with a 3-dimensional representation of (34) involving 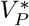, 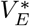, and 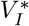, results in no loss of information. We therefore arrive at a final expression relating *Y*\* to external input-rate *P*, by substituting 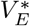 and 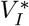 in the expression *Y* = *V_E_* = *V_I_*:

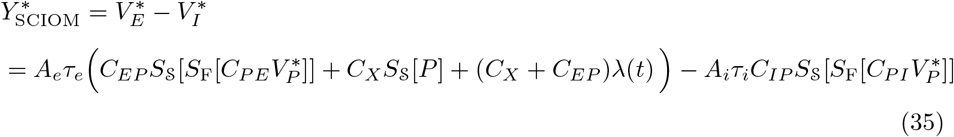

and then substituting 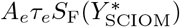 for 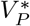, results in:

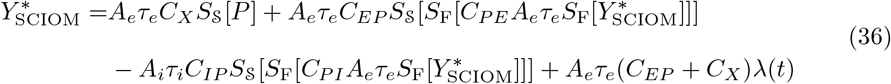

Equation (36) encapsulates the entire equilibria set of the SCIOM model. We are primarily interested in studying the family of equilibrium curves of the form 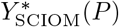 relating external input-rates *P* to critical states *Y*\*.

### Bifurcation Analysis of Curves *Y*\*(*P*)

This section outlines the bifurcation analysis used in this study to compare the family of equilibrium curves of the Jansen-Rit model, 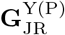, and the family of equilibrium curves of the SCIOM model, 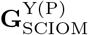. Analysis of these curves will describe exactly how the changes in the SCIOM model result in more physiologically realistic dynamics. These curves are acquired by manipulating the four intra-network coupling weights. Therefore, it is useful to define the space of intra-network coupling coefficients, **C**_W_:

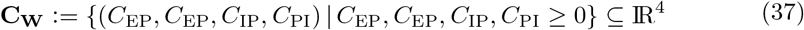

Now we can define the sets 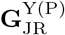 and 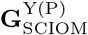 of equilibrium curves at different network coupling weights:

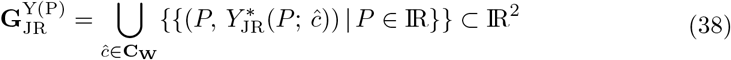

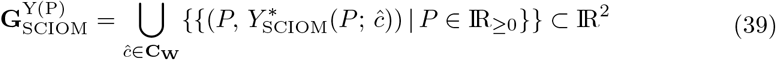

The rest of the parameters for each model are to be held fixed to the values defined in 1 and 2 for the respective model, unless otherwise specified.

More specifically, however, we are interested in subsets of 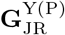 and 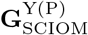 that satisfy certain criteria corresponding to dynamical regimes that are consistent with alpha-rhythm physiology in the visual cortex. We will discuss the criteria of these subsets and their corresponding dynamics. We will then introduce four equivalence relations over each equilibrium curve to outline the assortment of limit-set behaviors as a function of external drive *P*.

### Intrinsic Oscillations

To simulate the occurrence of alpha rhythms during quiescence we sought for a subset of curves 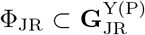 and 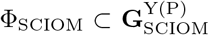 that possess an initial supercritical Hopf bifurcation at *P*_H_1__ ≤ 0 and a subsequent Hopf bifurcation somewhere between 0 ≤ *P*_H_2__ < 2*e_o_*. We also require for the decay of oscillatory power with increasing external drive. These three conditions can be written in a more formal manner:

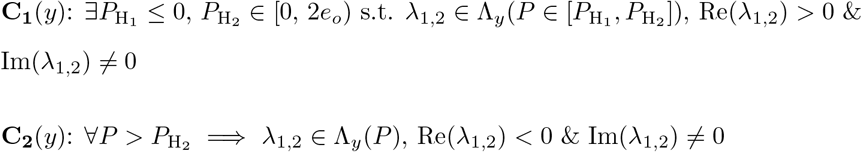

Where Λ_y_(*P*) is the diagonal matrix containing the eigenvalue pairs in the eigendcomposition of a system *y*’s Jacobian matrix at the specified point *P* (see Appendix C for more details).

Conditions **C**_1_ and **C**_2_ ensure the existence of stable oscillatory behavior in the absence of external drive. We refer to such oscillations as **intrinsic oscillations**. These two conditions also ensure the dissolution of stable oscillations within an external drive range of [0, 2*e_o_*). In addition to these conditions, we require a negative correlation between oscillatory power and external input *P*. Therefore, we enforce a third condition:

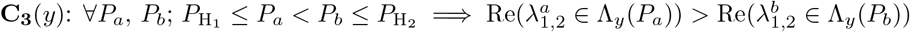

The set of equilibrium curves of the form *Y*\*(*P*) from the Jansen-Rit and SCIOM models that exhibit intrinsic oscillations are defined as such:

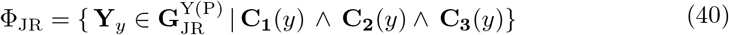

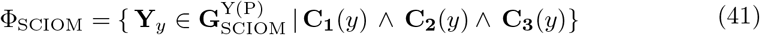

The sets Φ_JR_ and Φ_SCIOM_ correspond to parameter regimes that permit intrinsic oscillations for each model and that possess the ability to transmit a response transient.

### Partitions on *Y*\*(*P*)

We partition each curve in 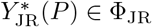 and 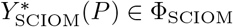 into four segments: the high and low fixed point segments Fix_H_ and Fix_L_, a separatrix, and the stable limit-cycle segment defined at its boundaries by an initial supercritical Hopf-bifurcation at one end and a subsequent Hopf-bifurcation at the other end. Examples of these curves with the following partitions indicated can be seen in 2. We’ll define these constructs over curves in Φ_SCIOM_, but their implications over curves in Φ_JR_ will be synonymous.

#### Fix_L_

The segment of fixed points 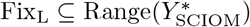 are stable states in terms of excitation-inhibition where the eigenvalues of the Jacobian matrix λ_*j*_ for each population *j* is Re(λ_*j*_) < 0 and Im(λ_*j*_) = 0.

#### Fix_H_

The segment of fixed points 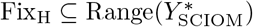 are stable states in terms of excitation-inhibition where the eigenvalues of the Jacobian matrix λ_*j*_ for each population *j* is *Re*(λ_*j*_) < 0 and Im(λ_*j*_) = 0. Although complex, the eigenpairs of the system y each have negative real parts, and so the system will converge to the fixed point.

#### Limit-cycle Segment (**LC**)

The segment of stable points in the 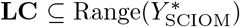 are stable states in terms of excitation-inhibition where the eigenvalues of the Jacobian matrix λ_*p*_ and λ*_i_* of the pyramidal and inhibitory populations each have *Re*(λ) > 0 and Im(λ) ≠ 0 parts. This results in the manifestation of stable oscillations overlying an unstable fixed point indicated by the positive real part of these eigenvalues.

#### Separatrix Region (**SP**)

The separatrix is a region in excitation-inhibition that exist between two stable segments. These are transient states that do not belong in the limit-set of the model.

### Simulations and Numerical Methods

#### Numerical Methods

Models were solved numerically using a 4th order Runge-Kutta integration algorithm in MatLAB 2019b (Mathworks. Natick, MA). Equations were integrated at 1kHz. XPP-AUT was used to calculate bifurcation points.

#### Phase Response Curves

Phase response curves were acquired by simulating an impulse stimulus to the SCIOM model at different phases of the oscillation between 0 and 2*π* and measuring phase change from the unperturbed case.

#### Coupling-Coefficient Surfaces

The 4-dimensional space of coupling weights **C**_W_ can be divided into two orthogonal half-spaces of 2-dimensions each: an **excitatory surface** *C*_EP_ × *C*_PE_ ⊂ **C**_W_, and an **inhibitory surface** *C*_IP_ × *C*_PI_ ⊂ **C**_W_. Within these half-spaces it is useful then to define regions that give rise to intrinsic oscillations. This region within the excitatory surface is acquired by fixing a point 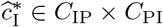, then,

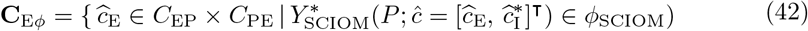

Likewise, a similar region in the inhibitory surface is acquired by fixing a point 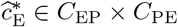, then,

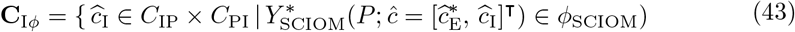

Depending on the choice of 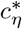, the orthogonal submanifold 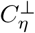 may be empty. The rest of the parameters of 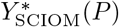 are held fixed to the values defined in 2, unless otherwise noted.

## Acknowledgements

We thank Audrey Sederberg for her helpful comments. This work was supported by National Institute of Health grant R01-MH118487.

## Appendices

## A Firing-rate to membrane potential operation

Each equation in system (18) serves to transform input-rates on the right-hand side to field-potentials on the left-hand side through the use of a conversion factor 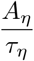. Where *τ_η_* determines the maximum synaptic response to a unit impulse function δ, and *τ_η_* determines its time course. To elucidate the synaptic response to *δ* we express the equation in terms of a second-order linear differential operator 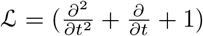:

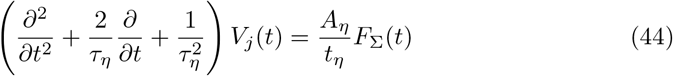

where *F*_Σ_(*t*) = *F_N_* (*t*) + *F_X_*(*t*) is the net input-rate to population *j*. In the case that *F*_Σ_(*t*) = *δ* is a dirac-delta function, *V_j_*(*t*) becomes an impulse response function *H_η_*:

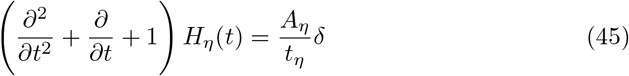

or more simply:

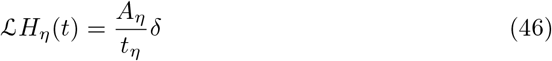

It can be inferred from (46) that the kernel of 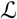, the impulse response function *H_η_*(*t*) must take the form:

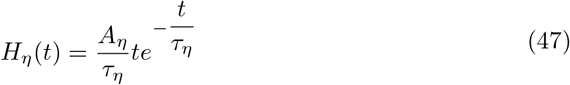

Therefore, the membrane potential of a population *j* at time *t* is the convolution of its impulse response function *H*_η_(*t*) with the net input-rate function *F*_Σ_(*t*) to the population:

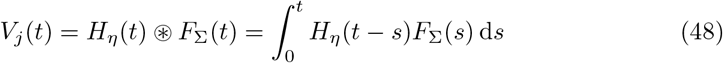

## C Linearization and Stability

The linearization of either system (JR or SCIOM) principally rests upon finding an ¦ approximation to *S*_F_ about the point 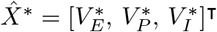. For the SCIOM system we ¦ consider the linearization of:

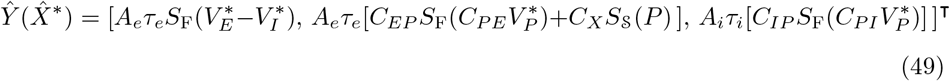

Its Jacobian **Dy**_*i,J*_(*x*), then takes the form:

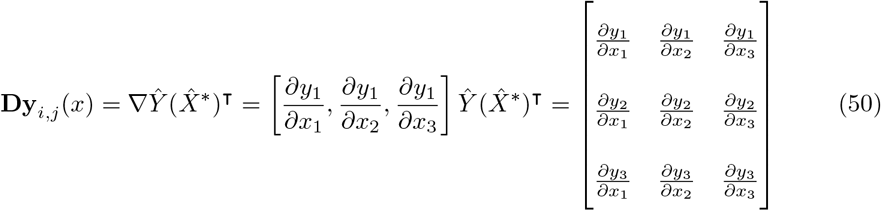

For stability and bifurcation analysis the eigendecomposition of its Jacobian matrix is considered at each point 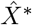,

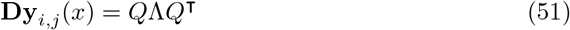

where *Q* is an orthogonal matrix of size 3 whose columns contain the corresponding eigenvectors of **Dy**_i,j_(*x*),

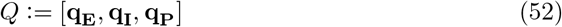

where **q**_E_, **q**_I_, **q**_P_ are the eigenvectors corresponding to the excitatory, pyramidal, and inhibitory components accordingly. The diagonal matrix of size *n*, Λ contains the eigenvalues of the Jacobian:

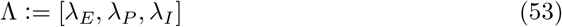

where λ_*E*_, λ_*P*_, and λ_*I*_ are the individual eigenvalues corresponding to the respective populations.

Of-course, because the nonlinearities of the model preclude analytical approaches, we used the numerical analysis software XPPAUT to detect both bifurcation points and stable regimes associated with the emergence and disappearance of oscillations within the domain of these curves.

## D Mapping between output Y and input P

The addition of a second nonlinearity, 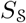, in the SCIOM model imposes several segments in which the system evolves independently of external input, driven only by the network interactions. In the Jansen-Rit paradigm the situation was reversed, the system would evolve in several segments independently of network interactions, driven solely by external inputs. However, in the SCIOM, the function 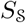 distorts the map of *P* to *Y*\* such that *P* ↦ *Y*\* is no longer a function, nor smooth at various points of *P*. We consider the closed-form expression of the map, *P* ↦ *Y*\*:

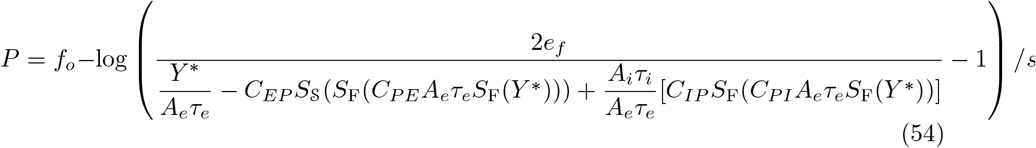

we can write this in a more concise and meaningful manner by defining:

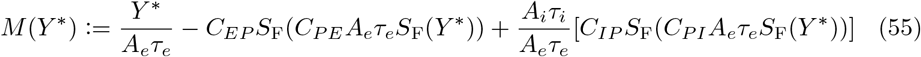

and noticing that *M* (*Y*\*) constitutes the synaptic drive to the system that evolves from external interactions. Put in terms of equation (29) for the synaptic drive *V_F_*(*t*), the variable 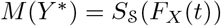. Now, we can re-write the expression (54) more concisely, using (55):

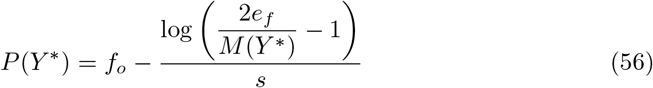

Equation (56) implies that the domain of synaptic drive, *M*(*Y*), influenced by external inputs is restricted between the bounds: 0 < *M*(*Y*\*) < 2*e_j_*, because *P*(*Y*) is undefined at 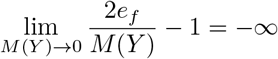 and as 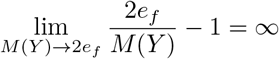. Synaptic drive outside of this dynamic range for external influences manifests from intranetwork interactions and from the relative internal state of the populations. Thus, in the SCIOM model, the system does change and responds with a signal Y in the absence of external influences at various points in its dynamics. Graphically, these points where Y changes independently of *P* fall within the vertical segments: **II**_1_, **II**_2_, and **II**_3_, illustrated in 2D.

On the contrary, the Jansen-Rit model is invariably influenced by external interactions. Even in domains where the network interactions are essentially shutoff or when the network is fully saturated. Notably, there are two half-lines that radiate away in a linear manner, mostly clearly see in 2A, from the dynamic core of the model. Here 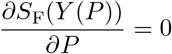, but the system continues to evolve as a function of *P* and with 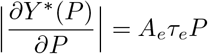.

The saturation point of *P* is not obvious when examining equation (55). In other words, it is not clear when increasing external input, *P*, begins to have negligible effect on synaptic drive. Since we run simulations at time steps of 0.001 seconds, we’ll consider a change of 0.001 in synaptic drive, *M*(*Y*), to be a least change. That is to say, 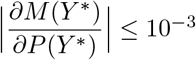 is consider negligible. Accordingly, *∂M*(*P*(*Y*\*) ≈10^3^) ≈8.982, therefore, ⌊*P*(*M*(*Y*\*) ≈8.982)⌋ = 17 Hz is the upper limit of external-input *P* in terms of generating a meaningful synaptic drive in the SCIOM model.

## E Excitatory/Inhibitory influence on state transitions: an energy and work analysis of network dynamics

From the equilibrium equation (25) of the Jansen-Rit model and (36) of the SCIOM model, it is apparent that the respective equilibria sets of the two models principally rest upon the weighted interactivity of the populations across the three firing-rate transformation functions in the system, *S*_f_. To study these interactions in more depth we define:

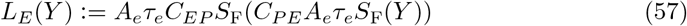

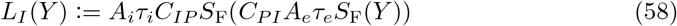

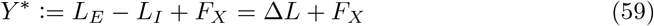

The function *L_E_* is the drive from the pyramidal population onto the excitatory population, and *L_I_* is the drive from the pyramidal population to the the inhibitory population.

To analyze these equations it is tempting to simplify them by replacing the sigmoid nonlinearty with a piecewise linear function or by reducing the dimensionality of the system through singular perturbation methods. However, it has been observed by Grimbert and Faugeras that these approximations destroy the oscillatory behavior. Therefore it appears that the oscillations emerging from the interactions between the excitation and inhibition loops is determined by higher order dynamics:

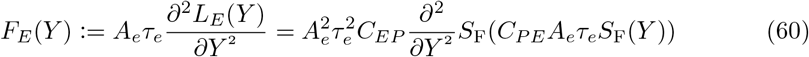

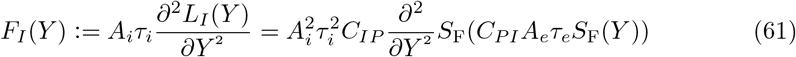

where 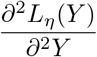 is weighted by the relevant post-synaptic potential amplitude and time constant, *A_η_τ_η_*. The function *F_E_* is the rate of change of the excitatory drive and *F_I_* measures rate of change of the inhibitory drive with respect to the model output, *Y*.

To better provide an intuitive analogy in understanding these equations, we employ an “energy function” to measure the stability across different output states of the system, *Y*:

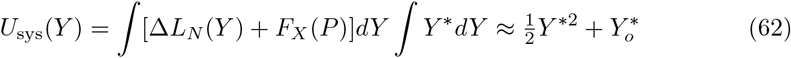

We refer to *U*_sys_(*Y*) as the potential-energy of the system at output *Y*. The constant of integration 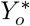 is used to offset the vertical shift in the graph of *Y*(*P*). Therefore, we take 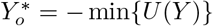, and sets our minimum value of *U*_sys_(*Y*) = 0.

Along with the potential-energy function *U*_sys_(*Y*), we define work done by the network using (60) and (61):

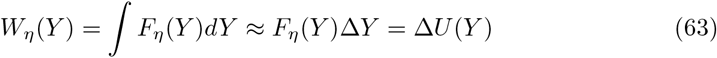

where we denote *W_η_* as the work done the loops within the network *L_η_*. More specifically, we are interested in the sum of the work done on the system by the excitation and inhibition loop interactions; thus,

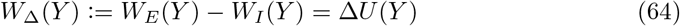

where *W*_Δ_(*Y*) is the difference between the work done on the system by the excitatory and inhibitory loop interactions which is equal to the change in the potential energy function of the system.

The amount of energy that is inserted into the system by the network interactions at output *Y* is calculated as:

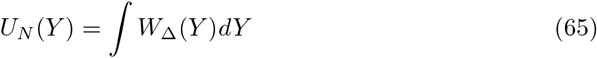

## References

[1] E D Adrian and B H C Matthews. “The berger rhythm: Potential changes from the occipital lobes in man”. In: Brain (1934). ISSN: 00068950. DOI: 10.1093/brain/57.4.355.

[2] Saeed Ahmadizadeh et al. “Bifurcation analysis of two coupled Jansen-Rit neural mass models.” In: PloS one 13.3 (2018). Ed. by Gennady Cymbalyuk, e0192842. ISSN: 1932-6203. DOI: 10.1371/journal.pone.0192842. URL: https://dx.plos.org/10.1371/journal.pone.0192842%20 http://www.ncbi.nlm.nih.gov/pubmed/29584728%20 http://www.pubmedcentral.nih.gov/articlerender.fcgi?artid=PMC5870965.

[3] Alan Anticevic and John D Murray. “Rebalancing Altered Computations: Considering the Role of Neural Excitation and Inhibition Balance Across the Psychiatric Spectrum.” In: Biological psychiatry 81.10 (2017), pp. 816–817. ISSN: 1873-2402. DOI: 10.1016/j.biopsych.2017.03.019. URL: http://www.ncbi.nlm.nih.gov/pubmed/28434614.

[4] K M Armstrong and T Moore. “{R}apid enhancement of visual cortical response discriminability by microstimulation of the frontal eye field”. In: Proc Natl Acad Sci U S A 104.22 (2007), pp. 9499–9504. DOI: 10.1073/pnas.0701104104.

[5] Thomas J Baumgarten, Alfons Schnitzler, and Joachim Lange. “Prestimulus Alpha Power Influences Tactile Temporal Perceptual Discrimination and Confidence in Decisions”. In: Cerebral Cortex 26.3 (Mar. 2016), pp. 891–903. ISSN: 14602199. DOI: 10.1093/cercor/bhu247. URL: http://www.ncbi.nlm.nih.gov/pubmed/25331603.

[6] H Berger. “Uber das Elektroenkephalogramm des Menschen. Arch. Pyschiatr. Nervenkr”. In: Arch. Pyschiatr. Nervenkr (1929).

[7] A Bollimunta et al. “Neuronal Mechanisms and Attentional Modulation of Corticothalamic Alpha Oscillations”. In: Journal of Neuroscience 31.13 (2011), pp. 4935–4943. ISSN: 0270-6474. DOI: 10.1523/JNEURUSCI.5580-10.2011. URL: http://www.jneurosci.org/cgi/doi/10.1523/JNEURUSCI.5580-10.2011.

[8] Anil Bollimunta et al. “Neuronal mechanisms of cortical alpha oscillations in awake-behaving macaques.” In: The Journal of neuroscience : the official journal of the S’ociety for Neuroscience 28.40 (2008), pp. 9976–9988. ISSN: 1529-2401. DOI: 10.1523/JNEURUSCI.2699-08.2008. URL: http://www.ncbi.nlm.nih.gov/pubmed/18829955%20 http://www.pubmedcentral.nih.gov/articlerender.fcgi?artid=PMC2692971.

[9] L J Borg-Graham, C Monier, and Y Frégnac. “Visual input evokes transient and strong shunting inhibition in visual cortical neurons.” In: Nature 393.6683 (May 1998), pp. 369–73. ISSN: 0028-0836. DOI: 10.1038/30735. URL: http://www.ncbi.nlm.nih.gov/pubmed/9620800.

[10] Michael E Brandt and Ben H Jansen. “The relationship between prestimulus alpha amplitude and visual evoked potential amplitude”. In: International Journal of Neuroscience 61.3-4 (1991), pp. 261–268. ISSN: 00207454. DOI: 10.3109/00207459108990744. URL: https://pubmed.ncbi.nlm.nih.gov/1824388/.

[11] Nicolas Brunel and Xiao-Jing Wang. “What determines the frequency of fast network oscillations with irregular neural discharges? I. Synaptic dynamics and excitation-inhibition balance.” In: Journal of neurophysiology 90.1 (July 2003), pp. 415–30. ISSN: 0022-3077. DOI: 10.1152/jn.01095.2002. URL: http://www.ncbi.nlm.nih.gov/pubmed/12611969.

[12] Daniel A Butts et al. “Temporal precision in the visual pathway through the interplay of excitation and stimulus-driven suppression.” In: The Journal of neuroscience : the official journal of the Society for Neuroscience 31.31 (Aug. 2011), pp. 11313–27. ISSN: 1529-2401. DOI: 10.1523/JNEUR0SCI.0434-11.2011. URL: http://www.ncbi.nlm.nih.gov/pubmed/21813691%20 http://www.pubmedcentral.nih.gov/articlerender.fcgi?artid=PMC3197857.

[13] Gyorgy Buzsaki,Costas A Anastassiou, and Christof Koch. “The origin of extracellular fields and currents-EEG, ECoG, LFP and spikes.” In: Nature reviews. Neuroscience 13.6 (2012), pp. 407–420. ISSN: 1471-0048. DOI: 10.1038/nrn3241. URL: https://www.nature.com/reviews/neuro%20 http://www.ncbi.nlm.nih.gov/pubmed/22595786%20 http://www.pubmedcentral.nih.gov/articlerender.fcgi?artid=PMC4907333.

[14] Aine Byrne, Daniele Avitabile, and Stephen Coombes. “Next-generation neural field model: The evolution of synchrony within patterns and waves”. In: Physical Review E 99.1 (2019). ISSN: 24700053. DOI: 10.1103/PhysRevE.99.012313.

[15] M H Chase and R M Harper. “Somatomotor and visceromotor correlates of operantly conditioned 12-14 c/sec sensorimotor cortical activity”. In: Electroencephalography and Clinical Neurophysiology (1971). ISSN: 00134694. DOI: 10.1016/0013-4694(71)90292-6.

[16] Vardhan S Dani et al. “Reduced cortical activity due to a shift in the balance between excitation and inhibition in a mouse model of Rett Syndrome”. In: Proceedings of the National Academy of Sciences of the United States of America 102.35 (2005), pp. 12560–12565. ISSN: 00278424. DOI: 10.1073/pnas.0506071102.

[17] Olivier David, Lee Harrison, and Karl J Friston. “Modelling event-related responses in the brain”. In: NeuroImage (2005). ISSN: 10538119. DOI: 10.1016/j.neuroimage.2004.12.030.

[18] Tolgay Ergenoglu et al. “Alpha rhythm of the EEG modulates visual detection performance in humans”. In: Cognitive Brain Resesarch (2004). ISSN: 09266410. DOI: 10.1016/j.cogbrainres.2004.03.009.

[19] Andreé A Fenton. “Excitation-inhibition discoordination in rodent models of mental disorders.” In: Biological psychiatry 77.12 (2015), pp. 1079–1088. ISSN: 1873-2402. DOI: 10.1016/j.biopsych.2015.03.013. URL: http://www.ncbi.nlm.nih.gov/pubmed/25895430%20 http://www.pubmedcentral.nih.gov/articlerender.fcgi?artid=PMC4444398.

[20] Alexander C Flint and Barry W Connors. “Two types of network oscillations in neocortex mediated by distinct glutamate receptor subtypes and neuronal populations”. In: Journal of Neurophysiology 75.2 (1996), pp. 951–957. ISSN: 0022-3077. DOI: 10.1152/jn.1996.75.2.951. URL: https://www.physiology.org/doi/10.1152/jn.1996.75.2.951.

[21] David C Godlove et al. “Microcircuitry of agranular frontal cortex: Testing the generality of the canonical cortical microcircuit”. In: Journal of Neuroscience (2014). ISSN: 15292401. DOI: 10.1523/JNEURUSCI.5127-13.2014.

[22] I C Gould,M F Rushworth, and A C Nobre. “Indexing the graded allocation of visuospatial attention using anticipatory alpha oscillations”. In: J Neurophysiol 105.3 (2011), pp. 1318–1326. DOI: 10.1152/jn.00653.2010.

[23] François Grimbert and Olivier Faugeras. “Analysis of Jansen’s model of a single cortical column”. In: Current Biology 14.1 (2006), p. 34. ISSN: 09609822. DOI: 10.1016/j.cub.2003.12.020. URL: https://hal.inria.fr/inria-00070410.

[24] Saskia Haegens et al. “α-Oscillations in the monkey sensorimotor network influence discrimination performance by rhythmical inhibition of neuronal spiking.” In: Proceedings of the National Academy of Sciences of the United States of America 108.48 (Nov. 2011), pp. 19377–82. ISSN: 1091-6490. DOI: 10.1073/pnas.1117190108. URL: http://www.ncbi.nlm.nih.gov/pubmed/22084106%20 http://www.pubmedcentral.nih.gov/articlerender.fcgi?artid=PMC3228466.

[25] Saskia Haegens et al. “Laminar Profile and Physiology of the α Rhythm in Primary Visual, Auditory, and Somatosensory Regions of Neocortex.” In: The Journal of neuroscience : the official journal of the Society for Neuroscience 35.42 (Oct. 2015), pp. 14341–52. ISSN: 1529-2401. DOI: 10.1523/JNEURUSCI.0600-15.2015. URL: http://www.jneurosci.org/content/35/42/14341.full.pdf%20 http://www.ncbi.nlm.nih.gov/pubmed/26490871%20 http://www.pubmedcentral.nih.gov/articlerender.fcgi?artid=PMC4683691.

[26] B Haider et al. “Neocortical network activity in vivo is generated through a dynamic balance of excitation and inhibition”. In: J Neurosci 26.17 (2006), pp. 4535–4545. DOI: 10.1523/JNEURUSCI.5297-05.2006.

[27] C C Hilgetag et al. “Anatomical connectivity defines the organization of clusters of cortical areas in the macaque monkey and the cat.” In: Philosophical transactions of the Royal Society of London. Series B, Biological sciences 355.1393 (Jan. 2000), pp. 91–110. ISSN: 0962-8436. DOI: 10.1098/rstb.2000.0551. URL: http://www.ncbi.nlm.nih.gov/pubmed/10703046%20 http://www.pubmedcentral.nih.gov/articlerender.fcgi?artid=PMC1692723.

[28] Luca Iemi et al. “Spontaneous Neural Oscillations Bias Perception by Modulating Baseline Excitability.” In: The Journal of neuroscience : the official journal of the S’ociety for Neuroscience 37.4 (2017), pp. 807–819. ISSN: 1529-2401. DOI: 10.1523/JNEURUSCI.1432-16.2016. URL: http://www.jneurosci.org/lookup/doi/10.1523/JNEUROSCI.1432-16.2017%20 http://www.ncbi.nlm.nih.gov/pubmed/28123017%20 http://www.pubmedcentral.nih.gov/articlerender.fcgi?artid=PMC6597018.

[29] Ben H Jansen and Michael E Brandt. “The effect of the phase of prestimulus alpha activity on the averaged visual evoked response”. In: Electroencephalography and Clinical Neurophysiology/Evoked Potentials Section 80.4 (1991), pp. 241–250. ISSN:01685597. DOI: 10.1016/0168-5597(91)90107-9. URL: https://linkinghub.elsevier.com/retrieve/pii/0168559791901079.

[30] Ben H Jansen and Vincent G Rit. “Electroencephalogram and visual evoked potential generation in a mathematical model of coupled cortical columns”. In: Biological Cybernetics 73.4 (1995), pp. 357–366. ISSN: 0340-1200. DOI: 10.1007/BF00199471. URL: http://link.springer.com/10.1007/BF00199471.

[31] Ben H Jansen, George Zouridakis, and Michael E Brandt. “A neurophysiologically-based mathematical model of flash visual evoked potentials”. In: Biological Cybernetics 68.3 (1993), pp. 275–283. ISSN: 03401200. DOI: 10.1007/BF00224863. URL: https://pubmed.ncbi.nlm.nih.gov/8452897/.

[32] Young Jin Jung, Kyung Hwan Kim, and Chang Hwan Im. Mathematical issues in the inference of causal interactions among multichannel neural signals. 2012. DOI: 10.1155/2012/472036.

[33] Colin Kehrer. “Altered excitatory-inhibitory balance in the NMDA-hypofunction model of schizophrenia”. In: Frontiers in Molecular Neuroscience 1 (2008), p. 6. ISSN: 1662-5099. DOI: 10.3389/neuro.02.006.2008.

[34] R C Kelly et al. “Local field potentials indicate network state and account for neuronal response variability”. In: J Comput Neurosci 29.3 (2010), pp. 567–579. DOI: 10.1007/s10827-009-0208-9.

[35] Wolfgang Klimesch, Robert Fellinger, and Roman Freunberger. “Alpha oscillations and early stages of visual encoding”. In: Front Psychol 2.MAY (2011), p. 118. ISSN: 1664-1078. DOI: 10.3389/fpsyg.2011.00118. URL: http://www.ncbi.nlm.nih.gov/pubmed/21687470%20 http://www.pubmedcentral.nih.gov/articlerender.fcgi?artid=PMC3108577%20 http://journal.frontiersin.org/article/10.3389/fpsyg.2011.00118/abstract.

[36] Wolfgang Klimesch, Paul Sauseng, and Simon Hanslmayr. EEG alpha oscillations: The inhibition-timing hypothesis. 2007. DOI: 10.1016/j.brainresrev.2006.06.003.

[37] Itamar D Landau et al. “The Impact of Structural Heterogeneity on Excitation-Inhibition Balance in Cortical Networks”. In: Neuron 92.5 (2016), pp. 1106–1121. ISSN: 10974199. DOI: 10.1016/j.neuron.2016.10.027.

[38] Katharina Limbach and Paul M Corballis. “Prestimulus alpha power influences response criterion in a detection task.” In: Psychophysiology 53.8 (2016), pp. 1154–64. ISSN: 1540-5958. DOI: 10.1111/psyp.12666. URL: http://www.ncbi.nlm.nih.gov/pubmed/27144476.

[39] K Linkenkaer-Hansen et al. “Prestimulus oscillations enhance psychophysical performance in humans”. In: J Neurosci 24.45 (2004), pp. 10186–10190. DOI: 10.1523/JNEURUSCI.2584-04.2004.

[40] M Lundqvist, P Herman, and A Lansner. “Effect of Prestimulus Alpha Power, Phase, and Synchronization on Stimulus Detection Rates in a Biophysical Attractor Network Model”. In: Journal of Neuroscience 33.29 (2013), pp. 11817–11824. ISSN: 0270-6474. DOI: 10.1523/JNEURUSCI.5155-12.2013. URL: http://www.ncbi.nlm.nih.gov/pubmed/23864671%20 http://www.pubmedcentral.nih.gov/articlerender.fcgi?artid=PMC3722510%20 http://www.jneurosci.org/cgi/doi/10.1523/JNEUROSCI.5155-12.2013.

[41] Daniel Malagarriga,Antonio J Pons, and Alessandro E P Villa. “Complex temporal patterns processing by a neural mass model of a cortical column.” In: Cognitive neurodynamics 13.4 (2019), pp. 379–392. ISSN: 1871-4080. DOI: 10.1007/s11571-019-09531-2. URL: http://www.ncbi.nlm.nih.gov/pubmed/31354883.

[42] Daniel Malagarriga et al. “Mesoscopic Segregation of Excitation and Inhibition in a Brain Network Model”. In: PLoS Computational Biology (2015). ISSN: 15537358. DOI: 10.1371/journal.pcbi.1004007.

[43] K E Mathewson et al. “Pulsed out of awareness: EEG alpha oscillations represent a pulsed-inhibition of ongoing cortical processing”. In: Front Psychol 2 (2011), p. 99. DOI: 10.3389/fpsyg.2011.00099.

[44] K E Mathewson et al. “To see or not to see: prestimulus alpha phase predicts visual awareness”. In: J Neurosci 29.9 (2009), pp. 2725–2732. ISSN: 1529-2401. DOI: 10.1523/JNEUR0SCI.3963-08.2009. URL: http://www.ncbi.nlm.nih.gov/pubmed/19261866.

[45] Carrie J McAdams and John H R Maunsell. “Effects of attention on orientation-tuning functions of single neurons in macaque cortical area V4”. In: Journal of Neuroscience (1999). ISSN: 02706474. DOI: 10.1523/jneurosci.19-01-00431.1999.

[46] T Moore and M Fallah. “Microstimulation of the frontal eye field and its effects on covert spatial attention”. In: J Neurophysiol 91.1 (2004), pp. 152–162. DOI: 10.1152/jn.00741.2002.

[47] Young MP et al. “Non-metric multidimensional scaling in the analysis of neuroanatomical connection data and the organization of the primate cortical visual system”. In: Philosophical transactions of the Royal Society of London. Series B, Biological sciences 348.1325 (1995), pp. 281–308. ISSN: 0962-8436. DOI: 10.1098/RSTB.1995.0069. URL: https://pubmed.ncbi.nlm.nih.gov/8577827/.

[48] B K Murphy and K D Miller. “Balanced amplification: a new mechanism of selective amplification of neural activity patterns”. In: Neuron 61.4 (2009), pp. 635–648. DOI: 10.1016/j.neuron.2009.02.005.

[49] John D Murray et al. “Linking Microcircuit Dysfunction to Cognitive Impairment: Effects of Disinhibition Associated with Schizophrenia in a Cortical Working Memory Model”. In: Cerebral Cortex 24.4 (2014), pp. 859–872. ISSN: 1047-3211. DOI: 10.1093/cercor/bhs370. URL: https://academic.oup.com/cercor/article-lookup/doi/10.1093/cercor/bhs370.

[50] Astrid A Prinz, Dirk Bucher, and Eve Marder. “Similar network activity from disparate circuit parameters.” In: Nature neuroscience 7.12 (Dec. 2004), pp. 1345–52. ISSN: 1097-6256. DOI: 10.1038/nn1352. URL: http://www.ncbi.nlm.nih.gov/pubmed/15558066.

[51] Rajasimhan Rajagovindan and Mingzhou Ding. “From prestimulus alpha oscillation to visual-evoked response: an inverted-U function and its attentional modulation.” In: Journal of cognitive neuroscience 23.6 (2011), pp. 1379–1394. ISSN: 0898-929X. DOI: 10.1162/jocn.2010.21478.

[52] Gustavo Rohenkohl and Anna C Nobre. “α oscillations related to anticipatory attention follow temporal expectations.” In: The Journal of neuroscience : the official journal of the Society for Neuroscience 31.40 (Oct. 2011), pp. 14076–84. ISSN: 1529-2401. DOI: 10.1523/JNEURUSCI.3387-11.2011. URL: http://www.ncbi.nlm.nih.gov/pubmed/21976492%20 http://www.pubmedcentral.nih.gov/articlerender.fcgi?artid=PMC4235253.

[53] A van Rotterdam et al. “A model of the spatial-temporal characteristics of the alpha rhythm”. In: Bulletin of Mathematical Biology 44.2 (1982), pp. 283–305. ISSN: 0092-8240. DOI: 10.1007/BF02463252. URL: http://link.springer.com/10.1007/BF02463252.

[54] J L R Rubenstein and M M Merzenich. Model of autism: Increased ratio of excitation/inhibition in key neural systems. 2003. DOI: 10.1034/j.1601-183X.2003.00037.x.

[55] Jason Samaha, Luca Iemi, and Bradley R Postle. “Prestimulus alpha-band power biases visual discrimination confidence, but not accuracy”. In: Consciousness and Cognition 54 (2017), pp. 47–55. ISSN: 10902376. DOI: 10.1016/j.concog.2017.02.005.

[56] Jason Samaha et al. “Top-down control of the phase of alpha-band oscillations as a mechanism for temporal prediction.” In: Proceedings of the National Academy of Sciences of the United States of America 112.27 (2015), pp. 8439–8444. ISSN: 1091-6490. DOI: 10.1073/pnas.1503686112. URL: http://www.ncbi.nlm.nih.gov/pubmed/26100913%20 http://www.pubmedcentral.nih.gov/articlerender.fcgi?artid=PMC4500260.

[57] T O Sharpee,K D Miller, and M P Stryker. “On the importance of static nonlinearity in estimating spatiotemporal neural filters with natural stimuli.” In: J Neurophysiol 99.5 (2008), pp. 2496–2509. URL: http://eutils.ncbi.nlm.nih.gov/entrez/eutils/elink.fcgi?cmd=prlinks&dbfrom=pubmed&retmode=ref&id=18353910.

[58] W L Shew et al. “Information capacity and transmission are maximized in balanced cortical networks with neuronal avalanches”. In: J Neurosci 31.1 (2011), pp. 55–63. DOI: 10.1523/JNEURUSCI.4637-10.2011.

[59] F H da Silva et al. “Model of brain rhythmic activity - The alpha-rhythm of the thalamus”. In: Kybernetik 15.1 (1974), pp. 27–37. ISSN: 03401200. DOI: 10.1007/BF00270757.

[60] R M Smeal,G B Ermentrout, and J A White. “Phase-response curves and synchronized neural networks”. In: Philos Trans R Soc Lond B Biol Sci 365.1551 (2010), pp. 2407–2422. DOI: 10.1098/rstb.2009.0292.

[61] Roberto C Sotero et al. “Realistically coupled neural mass models can generate EEG rhythms”. In: Neural Computation (2007). ISSN: 1530888X. DOI: 10.1162/neco.2007.19.2.478.

[62] Eelke Spaak et al. “Layer-specific entrainment of γ-band neural activity by the α rhythm in monkey visual cortex.” In: Current biology : CB 22.24 (Dec. 2012), pp. 2313–8. ISSN: 1879-0445. DOI: 10.1016/j.cub.2012.10.020. URL: http://www.ncbi.nlm.nih.gov/pubmed/23159599%20 http://www.pubmedcentral.nih.gov/articlerender.fcgi?artid=PMC3528834.

[63] P Suffczynski et al. “Computational model of thalamo-cortical networks: dynamical control of alpha rhythms in relation to focal attention”. In: Int J Psychophysiol 43.1 (2001), pp. 25–40. DOI: 10.1016/s0167-8760(01)00177-5.

[64] G Thut et al. “Alpha-band electroencephalographic activity over occipital cortex indexes visuospatial attention bias and predicts visual target detection”. In: J Neurosci 26.37 (2006), pp. 9494–9502. DOI: 10.1523/JNEURUSCI.0875-06.2006.

[65] Gina G. Turrigiano et al. Rapid and Active Stabilization of Visual Cortical Firing Rates Across Light-Dark Transitions. 2019. DOI: 10.1101/542670.

[66] Fabrice Wendling et al. “Epileptic fast activity can be explained by a model of impaired GABAergic dendritic inhibition”. In: European Journal of Neuroscience 15.9 (2002), pp. 1499–1508. ISSN: 0953816X. DOI: 10.1046/j.1460-9568.2002.01985.x.

[67] Hugh R Wilson and Jack D Cowan. “Excitatory and inhibitory interactions in localized populations of model neurons.” In: Biophysical journal 12.1 (1972), pp. 1–24. ISSN: 0006-3495. DOI: 10.1016/S0006-3495(72)86068-5. URL: http://www.ncbi.nlm.nih.gov/pubmed/4332108%20 http://www.pubmedcentral.nih.gov/articlerender.fcgi?artid=PMC1484078.

[68] Mingshan Xue, Bassam V Atallah, and Massimo Scanziani. “Equalizing excitation-inhibition ratios across visual cortical neurons”. In: Nature 511.7511 (2014), pp. 596–600. ISSN: 14764687. DOI: 10.1038/nature13321.

[69] Ofer Yizhar et al. “Neocortical excitation/inhibition balance in information processing and social dysfunction”. In: Nature 477.7363 (2011), pp. 171–178. ISSN: 00280836. DOI: 10.1038/nature10360. URL: https://www.nature.com/articles/nature10360.

[70] Edward Zagha, John D Murray, and David A McCormick. “Simulating Cortical Feedback Modulation as Changes in Excitation and Inhibition in a Cortical Circuit Model”. In: eneuro 3.4 (2016), pp. 208–216. ISSN: 2373-2822. DOI: 10.1523/ENEURU.0208-16.2016. URL: http://eneuro.org/lookup/doi/10.1523/ENEURU.0208-16.2016.

